# A temporal threshold in TGF-β signaling separates reversible tissue adaptation from persistent NK cell effector dysfunction

**DOI:** 10.64898/2026.02.11.705354

**Authors:** Kevin Schmid, Christina Haimerl, Julia Stark, Erik Müller, Leon Merz, Robin Schenk, Karyna Pistrenko, Julia Baumgarten, Kim Berberich, Ulrike Bauer, Ursula Ehmer, Melanie Laschinger, Norbert Hüser, Daniel Hartmann, Jan P. Böttcher, Roland M. Schmid, Gabriela M. Wiedemann

**Affiliations:** TUM School of Medicine and Health - Clinical Department of Internal Medicine II, TUM University Hospital; Munich, Germany; TUM School of Medicine and Health - Institute of Molecular Immunology, TUM University Hospital; Munich, Germany; TUM School of Medicine and Health - Department of Surgery, TUM University Hospital; Munich, Germany; Department of General, Visceral and Transplantation Surgery, University Hospital Tübingen, Hoppe-Seyler-Strasse 3, 72076; Tübingen, Germany; Department of Experimental Immunology, Institute of Immunology, Eberhard Karls University of Tübingen; Tübingen, Germany; M3 Research Center, University Hospital Tübingen, Eberhard Karls University of Tübingen; Tübingen, Germany; Cluster of Excellence iFIT (EXC 2180) ’Image Guided and Functionally Instructed Tumor Therapies’, Eberhard Karls University of Tübingen; Tübingen, Germany

## Abstract

TGF-β signaling is a major regulator of immune cell differentiation and function, yet whether prolonged signaling can durably imprint dysfunctional immune states independently of continued pathway engagement remains unclear. Here, we identify signal duration as a critical determinant of human natural killer (NK) cell fate: short-term TGF-β exposure induces largely reversible transcriptional, chromatin and functional changes, whereas prolonged exposure establishes persistent effector dysfunction independent of continued signaling. Mechanistically, prolonged TGF-β drives durable loss of chromatin accessibility at effector-associated regulatory elements, particularly those enriched for IRF, EOMES, and T-bet binding motifs, resulting in stable restriction of NK cell effector programs despite reversibility of active histone marks. In contrast, tissue residency-associated programs remain largely reversible, revealing that distinct NK cell fate programs differ fundamentally in their susceptibility to epigenetic fixation downstream of the same cytokine signal. SMAD4 CUT&RUN and knockout experiments further demonstrate that canonical TGF-β signaling acts primarily through rewiring of upstream transcriptional regulatory networks rather than direct targeting of effector loci. Finally, NK cells from patients with hepatocellular carcinoma recapitulate key functional and epigenetic features of the persistent TGF-β-associated state defined *in vitro*. Together, these findings identify signal duration as a critical determinant of TGF-β-driven cell fate and demonstrate that prolonged TGF-β exposure can induce epigenetically stabilized dysfunctional states that persist after signal withdrawal, with important implications for therapeutic strategies aimed at reversing chronic TGF-β-mediated dysfunction.

## INTRODUCTION

Transforming growth factor-β (TGF-β) is a major regulator of immune cell differentiation, tissue adaptation, and suppression of anti-tumor immunity^1^. A key unresolved question is whether prolonged TGF-β signaling can epigenetically fix lymphocyte dysfunction independently of continued pathway activity. This distinction has important implications for understanding chronic immune dysfunction in cancer and for therapeutic strategies aimed at reversing TGF-β-driven suppression.

Natural killer (NK) cells are a compelling system in which to study this question because their functional state is highly responsive to environmental cytokine cues and they can undergo durable epigenetic remodeling following activation^2, 3^. NK cells are innate lymphocytes that rapidly eliminate tumor cells in an antigen-independent manner and shape adaptive immunity through cytokine and chemokine secretion, positioning them as powerful effectors for cancer immunotherapy ^4, 5, 6, 7, 8^. In solid tumors, however, NK cell function is profoundly curtailed by immunosuppressive cues within the tumor microenvironment, and restoring it remains a central therapeutic challenge ^7, 9, 10^. Among these inhibitory signals, TGF-β stands out as a major suppressor diverse effector programs, dampening cytotoxicity, cytokine production, and cellular metabolism, and promoting conversion toward a tissue-resident ILC1-like phenotype with reduced anti-tumor activity ^11, 12, 13, 14, 15, 16, 17, 18^.

A critical unresolved question is whether TGF-β-mediated NK cell dysfunction remains continuously dependent on suppressive signaling or instead becomes epigenetically fixed. Although TGF-β has been linked to transcriptional and epigenetic remodeling in dysfunctional NK cells^19, 20^, whether these changes persist after signal withdrawal, which functional programs become durably stabilized, and whether signal duration itself determines the permanence of suppression remain unknown.

That cytokine signals can durably encode NK cell fate is well established in the context of activating stimuli ^2, 3, 21, 22, 23^. Cytokine-induced memory-like (CIML) NK cells generated by combined exposure to IL-12, IL-15, and IL-18 acquire stably enhanced effector function through persistent epigenetic remodeling that remains detectable long after cytokine withdrawal ^21, 24^.

Whether suppressive cytokine signaling can similarly impose durable dysfunctional states has not been established. Canonical TGF-β signaling proceeds through SMAD2/3-SMAD4 complexes that cooperate with partner transcription factors (TFs) to regulate transcriptional programs and associated chromatin states at target loci, while non-canonical pathways contribute additional layers of context-dependent regulation ^25, 26^. Whether these signaling programs can drive stable epigenetic fixation of NK cell dysfunction remains unknown.

Here we show that the duration of TGF-β exposure determines the stability of human NK cell dysfunction. We demonstrate that short-term TGF-β exposure induces largely reversible transcriptional and chromatin changes, whereas prolonged exposure drives durable remodeling of effector-associated regulatory elements, marked by loss of accessibility at IRF, EOMES, and T-bet binding sites, that persists after signal withdrawal. In contrast, tissue residency (TR) genes escape this epigenetic fixation, revealing that effector programs and tissue adaptation are governed by distinct regulatory mechanisms downstream of TGF-β. NK cells from hepatocellular carcinoma (HCC) patients display impaired effector function and chromatin alterations overlapping with those induced by prolonged TGF-β exposure, consistent with chronic *in vivo* signaling as a driver of persistent NK cell dysfunction in human cancer. SMAD4 perturbation and CUT&RUN analyses further indicate that canonical TGF-β signaling stabilizes dysfunction primarily through rewiring of upstream transcriptional networks rather than direct silencing of effector loci. Together, these findings identify TGF-β signal duration as a determinant of NK cell fate and establish persistent NK cell dysfunction as an epigenetically stabilized state, with broader implications for understanding how chronic suppressive signaling shapes durable immune cell dysfunction.

## RESULTS

### TGF-β signal duration governs transcriptional and epigenetic outcomes in NK cells

To determine whether short-term and prolonged TGF-β exposure engage distinct gene regulatory programs, we performed RNA-sequencing (RNA-seq) and assay for transposase-accessible chromatin with sequencing (ATAC-seq) on sorted human NK cells (CD3⁻CD14⁻CD19⁻CD56⁺) following short-term (3 hours, a timepoint at which canonical SMAD2/3 phosphorylation is already robustly induced) or prolonged (7 or 17 days) TGF-β exposure (**Fig. S1 A-C**). After 3 hours, transcriptional changes were modest with 427 differentially expressed genes (DEGs) in total, and dominated by rapid upregulation of negative regulators of TGF-β signaling including SMAD7, SMURF2, and SKI, with limited effects on effector or TR genes (**Fig. 1A, Fig. S1D**). By contrast, prolonged stimulation drove extensive transcriptional reprogramming: 7-day exposure resulted in 966 DEGs, with downregulation of key effector genes including *IFNG*, *GZMA*, and *PRF1*, alongside upregulation of TR markers such as *ITGAE* and *ITGA1*, a pattern further consolidated after 17 days (**Fig. 1A, Fig. S1D**). At the epigenetic level, even brief TGF-β exposure induced chromatin remodeling with 2,314 differentially accessible regions (DARs) predominantly at intergenic and intronic putative enhancer elements, but prolonged exposure markedly amplified this response (26,538 DARs after 7 days and 27,802 DARs after 17 days of TGF-β) and shifted the balance toward chromatin closure (**Fig. 1B, Fig. S1D**).

**Figure 1.**
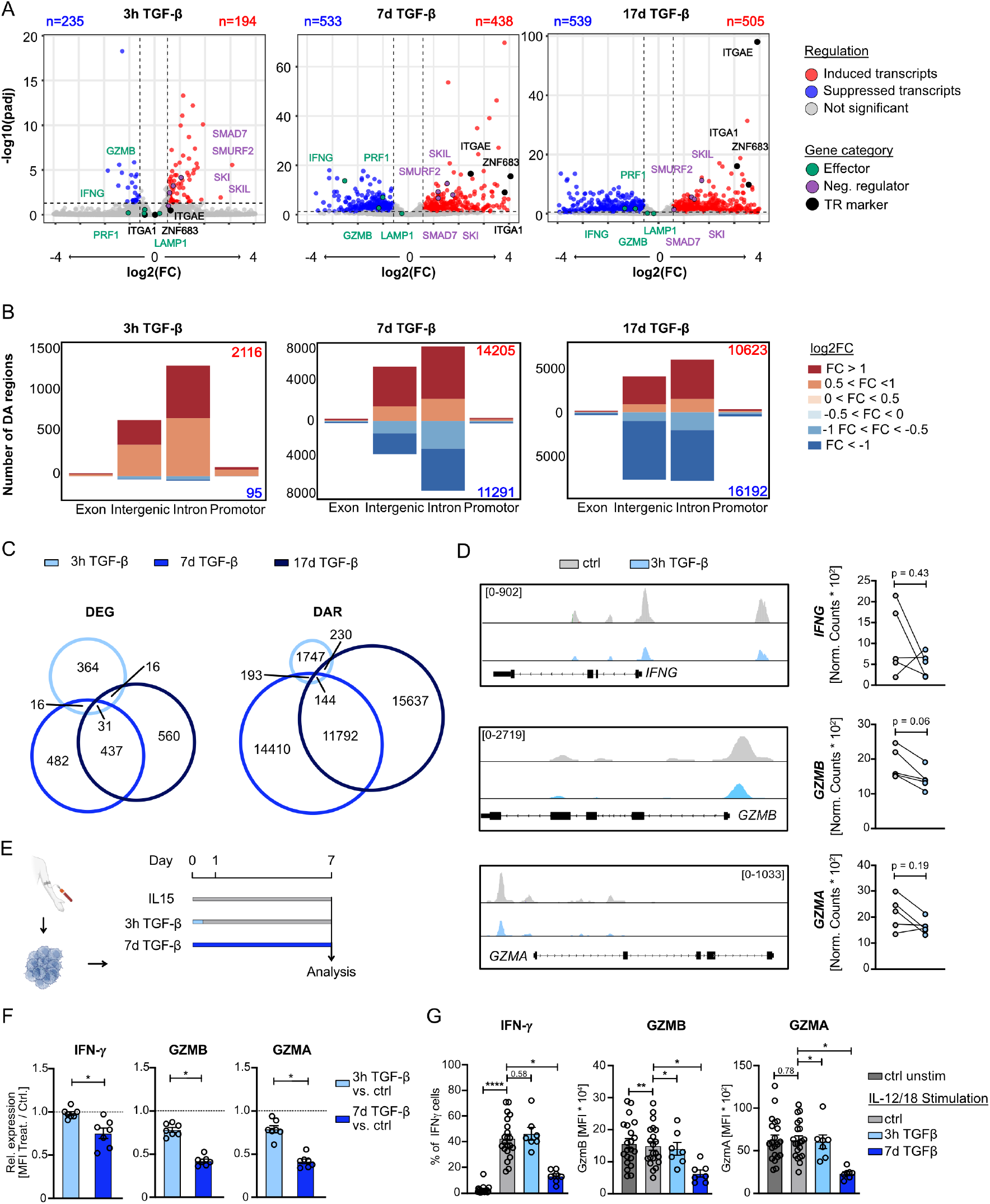
TGF-β signal duration shapes transcriptional and epigenetic responses. Human NK cells were sorted and treated with TGF-β [10 ng/mL] and IL-15 [50 ng/mL] for the indicated durations, followed by transcriptomic and epigenomic profiling. **(A)** Volcano plots showing DEGs between control and 3 hours (h), 7 days (d) or 17 d treated NK cells (padj < 0.05). **(B)** Peak type analysis of differentially accessible (DA) peaks in ATAC seq. Comparison of 3 h, 7 d and 17 d TGF-β treated NK cells versus control. **(C)** Venn diagram comparison of DEGs and DARs. **(D)** Genome browser (IGV) tracks showing chromatin accessibility at representative effector loci in control, 3 h TGF-β-treated NK cells. Tracks are counts per million (CPM) normalized. Paired dot plot depicting normalized counts for promoter regions of selected genes. **(E)** Experimental setup in which NK cells were treated with IL-15 only (control; 50 ng/mL), with TGF-β for 3 hours followed by washing and return to IL-15-only culture for an additional 7 d, or with TGF-β plus IL-15 for 7 d. All analyses were performed on d 7. **(F)** Ratio of mean fluorescence intensity (MFI) of IFN-γ and GZMA in TGF-β-treated versus control NK cells at day 7 (mean ± SEM; n = 7; two independent experiments). **(G)** Functional response of NK cells restimulated with IL-12 (20 ng/mL) and IL-18 (10 ng/mL) at d 7. Bar plots show frequency or MFI; control conditions are pooled for each time point (n = 7; two independent experiments). Significance was calculated using a Wilcoxon signed-rank test (*p<0.05; **p<0.01; ***p<0.001; **p<0.0001).

Gene set enrichment analysis (GSEA) revealed a duration-dependent shift in the transcriptional consequences of TGF-β signaling. Whereas short-term exposure primarily induced canonical TGF-β-associated pathways, prolonged stimulation was characterized by progressive suppression of oxidative phosphorylation, interferon response, and MYC target programs (**Fig. S1E**).

Comparison of DEGs across timepoints revealed limited overlap between short- and long-term transcriptional responses. The small shared gene set of 31 genes was enriched for negative regulators of TGF-β signaling - including *SMAD7, SMURF2*, and *SKI* - suggesting that feedback control represents a conserved transcriptional response independent of signal duration (**Fig. 1C**). Similarly, short- and long-term chromatin accessibility programs showed minimal overlap, indicating that extensive epigenetic remodeling requires prolonged TGF-β exposure (**Fig. 1C**).

Among the shared DARs, increased accessibility at the *SMAD7* locus reinforced the negative feedback response, while decreased accessibility at regions associated with *STAT4* and *NFATC2* suggested concomitant suppression of key activating TFs.

Of note, even short-term TGF-β exposure showed a trend toward decreased accessibility at key effector loci, including IFNG, GZMA, and GZMB in a subset of donors (**Fig. 1D**), suggesting that epigenetic remodeling at critical effector genes may begin early during TGF-β signaling.

To address whether this early epigenetic remodeling is sufficient to induce lasting functional changes, NK cells were stimulated with TGF-β for 3 h, followed by washout and a 6-day resting period, and compared to NK cells continuously exposed to TGF-β for 7 days as a model of chronic signaling. Prolonged TGF-β exposure induced robust reductions in granzyme A, granzyme B, and IFN-γ expression, whereas 3 h exposure with subsequent withdrawal resulted in only minimal changes (**Fig. 1F**). Critically, when subsequently challenged with IL-12/IL-18 restimulation, NK cells exposed to TGF-β for 3 h and rested for 6 days retained largely normal cytokine responsiveness, whereas those exposed for 7 days showed markedly blunted IFN-γ production and granzyme expression even under activating conditions (**Fig. 1G**), indicating that short-term TGF-β exposure is insufficient to induce sustained dysfunction. Collectively, these data demonstrate that TGF-β reprograms NK cells in a duration-dependent manner, with short-term exposure initiating limited and reversible changes, while prolonged signaling drives extensive epigenetic remodeling, raising the question of whether these effects are transient or give rise to lasting NK cell dysfunction.

### Effector dysfunction but not TR programming persists after TGF-β withdrawal

We next asked whether prolonged TGF-β signaling induces a persistent dysfunctional state that remains after signal withdrawal. To address this, NK cells were stimulated with TGF-β for 7 days, followed by a 10-day resting period, and subsequently analyzed for effector function, metabolic fitness, and expression of TR-associated markers (**Fig. 2A**). Sustained TGF-β exposure (17 days) robustly induced TR-associated markers CD103, CD9, and CD49a, while reducing intracellular granzyme A, granzyme B, and IFN-γ levels (**Fig. 2B-C**). Following TGF-β withdrawal for 10 days, TR markers largely returned to baseline, underscoring the requirement for continuous TGF-β signaling to maintain this phenotype (**Fig 2B**).

**Figure 2.**
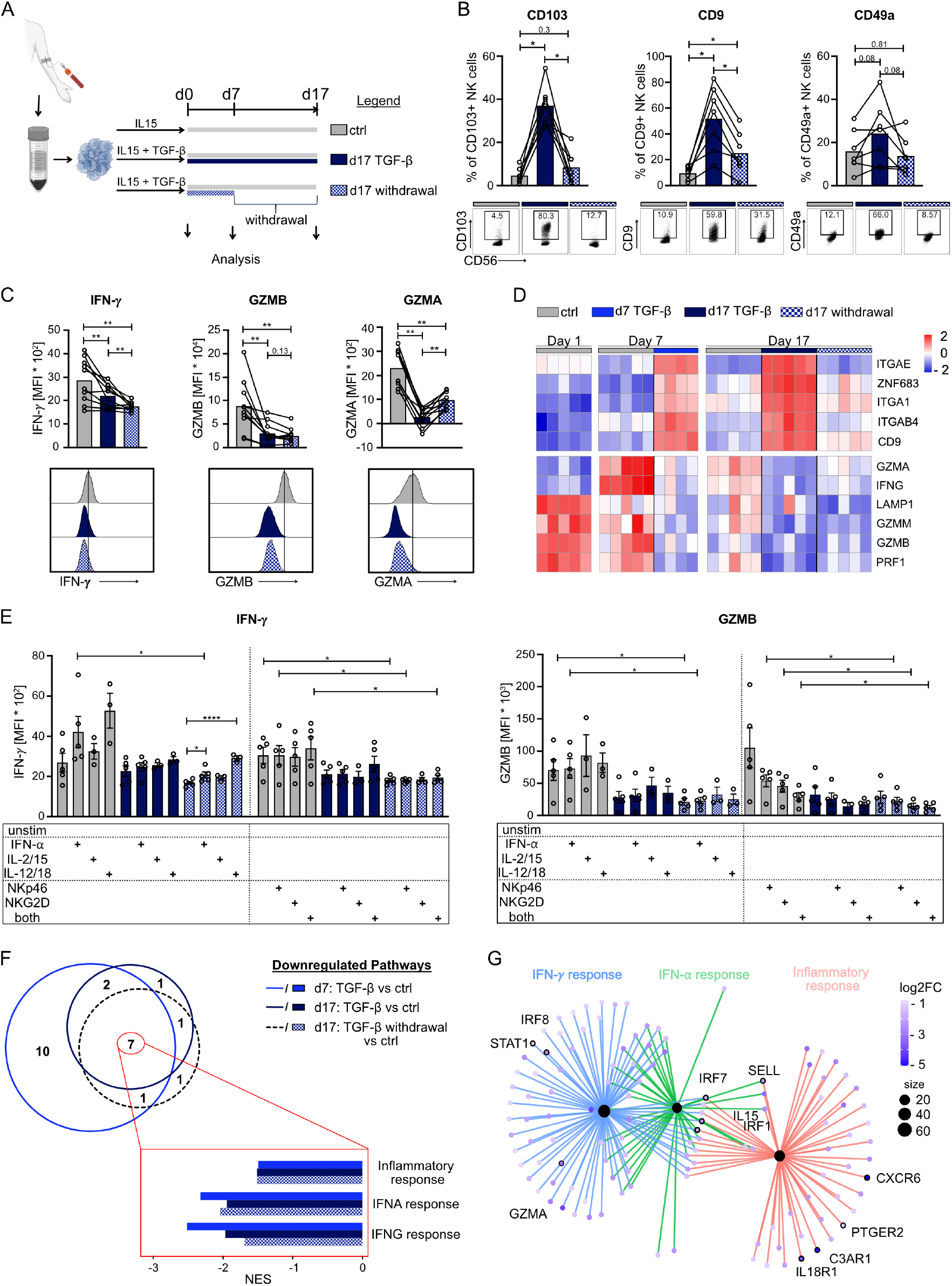
Prolonged TGF-β signaling induces durable impairment of the NK cell effector program. **(A)** Experimental design. Enriched primary human NK cells were cultured for 7 d in IL-15 [50 ng/mL] with or without TGF-β [10 ng/mL]. On day 7, media were exchanged for all conditions, and cells were maintained in IL-15 (control) or IL-15 + TGF-β. For the withdrawal condition, cells were washed twice on d 7 and cultured in IL-15 for an additional 10 d. Analyses were performed at days 0, 7, and 17. **(B)** Baseline frequencies of CD103⁺, CD9⁺, and CD49a⁺ NK cells at d 17, shown as bar plots (n = 7; three independent experiments). A representative donor is shown as a dot plot. **(C)** Baseline frequencies of granzyme A-positive (GZMA⁺), granzyme B-positive (GZMB⁺), and IFN-γ⁺ NK cells at day 17, assessed by flow cytometry and shown as bar plots (n = 7; three independent experiments). A representative donor is shown as a histogram. **(D)** RNA-seq analysis at the indicated time points. Heatmap shows z-score of regularized counts of selected TR-associated and effector genes across individual donors (n = 5). **(E)** NK cells were restimulated at day 17 as indicated and analyzed by flow cytometry. Bar plots show mean fluorescence intensity (MFI) of GZMA and IFN-γ (n = 5; three independent experiments). A representative donor is shown as a histogram. Statistical significance was calculated only between control and withdrawal conditions and within the withdrawal group; only significant comparisons are shown. **(F)** Venn diagram showing overlap of downregulated gene sets (Hallmark database; NES < 0, padj < 0.05) across the indicated conditions (n = 5). Vertical bar graphs depict normalized enrichment scores (NES) for selected effector-associated gene sets. **(G)** Network representation of connectivity between selected pathways (cnetplot; clusterProfiler), with genes shown as nodes colored by log2(FC) expression values. Across the figure, statistical significance was determined using a Wilcoxon signed-rank test for panels B and C and a paired Student’s t test for panel E (*p < 0.05; **p < 0.01; ***p < 0.001; ****p < 0.0001).

To exclude the possibility that loss of TR features reflected selective expansion of TR-negative cells, CD103⁺ and CD103⁻ NK cells were sorted after 7 days of TGF-β exposure and cultured separately (**Fig. S2A-B**). Both populations showed comparable loss of CD103 expression following TGF-β withdrawal (**Fig. S2C**), demonstrating that even CD103⁺ NK cells require sustained signaling to maintain TR marker expression. Moreover, effector dysfunction was maintained independently of CD103 expression status, further supporting a dissociation between TR-associated features and persistent functional impairment (**Fig. S2C**).

In contrast to the reversibility of TR features, NK cell effector function demonstrated sustained impairment after TGF-β withdrawal. Reduced expression of granzyme A and B and IFN-γ was maintained even after 10 days of rest (**Fig. 2C**), indicating long-lasting functional suppression independent of ongoing signaling. This suppression was not attributable to metabolic defects, as mitochondrial mass, and mitochondrial membrane potential were comparable between TGF-β-exposed and control NK cells at day 17 (**Fig. S2D**).

Consistent with these findings, RNA-seq analysis across all time points revealed that TR-associated transcripts (*ITGAE*, *ZNF683*, *ITGA1*, *ITGB4*, *CD9*) were selectively upregulated only under continuous TGF-β exposure, but returned to baseline after withdrawal (**Fig. 2D**). In contrast, effector genes (*GZMA*, *GZMB*, *PRF1*, *LAMP1*, *IFNG*) remained persistently downregulated even after prolonged signal withdrawal (**Fig. 2D**). NK cell proliferation was likewise durably impaired, as reflected by reduced Ki67 expression (**Fig. S2E**).

To determine whether this defect reflected disruption of specific activation pathways, TGF-β-naïve and TGF-β-exposed NK cells (with or without recovery) were restimulated using diverse activating cues, including IFN-α, IL-2 plus IL-15, IL-12 plus IL-18, and activating receptor ligation. Intriguingly, none of these stimuli fully restored NK cell effector function (**Fig. 2E, S2F**), indicating broad functional impairment spanning multiple activating pathways. Consistent with this broad functional impairment, GSEA identified inflammatory response as well as IFN-α and IFN-γ signaling pathways among the most persistently suppressed programs across all prolonged exposure conditions, including after withdrawal (**Fig. 2F**). These pathways encompassed key TFs such as *STAT1*, *IRF1*, and *IRF8*, along with effector-associated genes like *GZMA*, and *IL18R1* (**Fig. 2G**). Together, these findings reveal a striking divergence in the stability of TGF-β-induced NK cell programs: whereas TR-associated features remain dependent on continuous signaling, effector dysfunction persists independently of cytokine withdrawal.

### Functional impairment after prolonged TGF-β exposure is accompanied by persistent epigenetic reprogramming

Given the durable functional impairment observed after prolonged TGF-β exposure, we next asked whether stable epigenetic programs underlie the persistent NK cell phenotype. To test this, we performed RNA-seq and ATAC-seq on day 17, comparing TGF-β-naïve NK cells, NK cells continuously exposed to TGF-β for 17 days, and NK cells exposed for 7 days followed by 10 days of signal withdrawal. To focus specifically on TGF-β-driven effects, we restricted our analysis to genes whose regulation at day 7 was attributable to TGF-β rather than IL-15 **(Fig. S3A**).

Within this TGF-β-regulated gene set, 1,071 DEGs were identified on day 17 in continuously treated NK cells, while 228 DEGs remained differentially expressed after TGF-β withdrawal, compared to controls (**Fig. 3A**). Temporal clustering revealed four distinct expression patterns reflecting transient or persistent up- or downregulation (**Fig. 3B**). Transiently upregulated genes that returned to baseline after withdrawal included TIMP1 and CTLA4, whereas persistently upregulated genes included SMAD3. In contrast, transiently downregulated genes included KLF2, consistent with the reversibility of TR features, while persistently downregulated genes comprised key effector molecules such as GZMB, PRF1, and LAMP1 (**Fig. 3B**).

**Figure 3.**
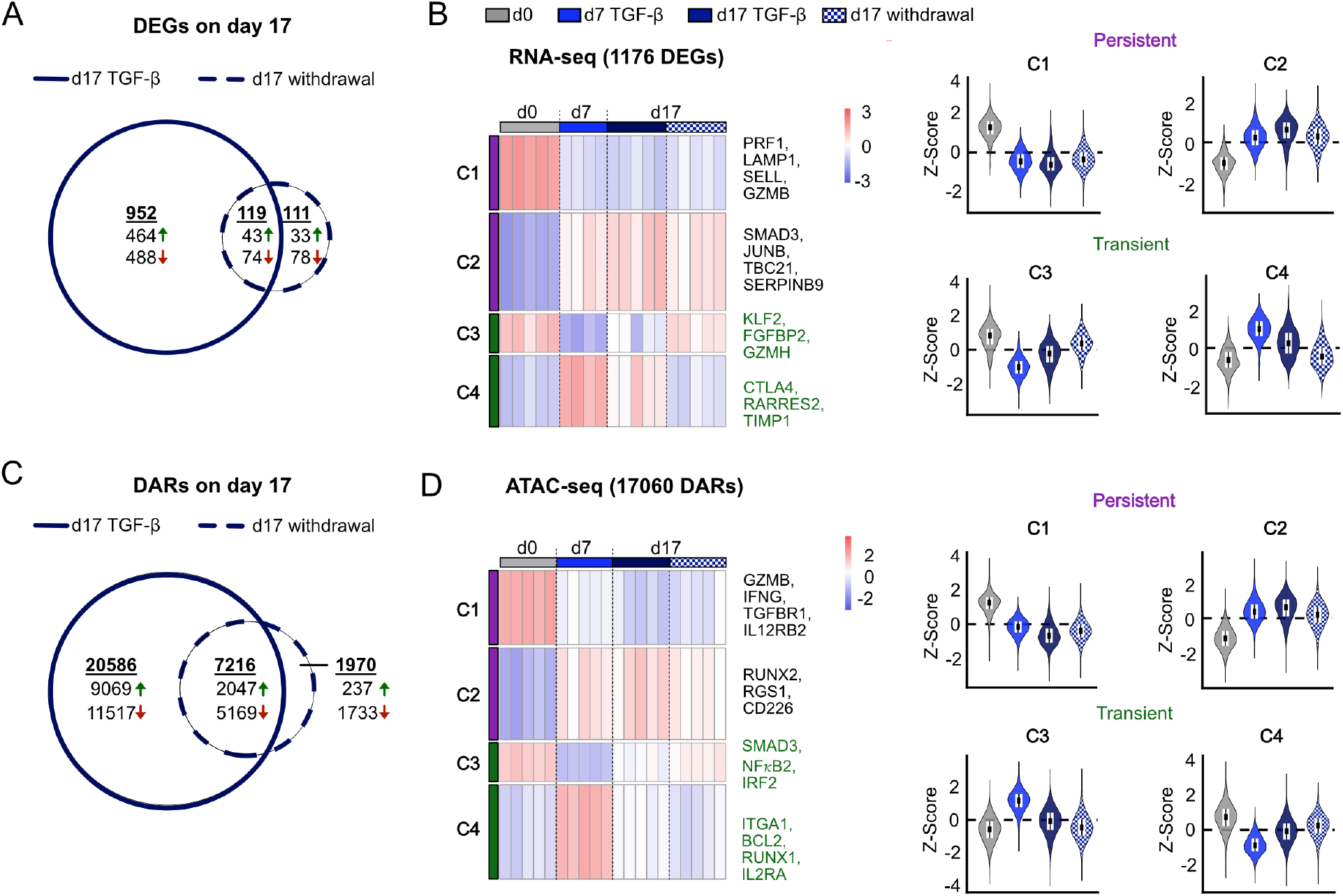
Prolonged TGF-β signaling drives lasting epigenetic changes. Primary human NK cells were cultured for 7 d in IL-15 with or without TGF-β. On d 7, media were exchanged for all conditions, and cells were maintained in fresh IL-15 (control) or IL-15 + TGF-β. For the withdrawal condition, cells were washed twice on d 7 and cultured in IL-15 for an additional 10 days. RNA-seq and ATAC-seq were performed at days 0, d 7, and d 17. **(A)** Euler diagram showing the overlap of differentially expressed genes (DEGs; |log2(FC)| > 0.58, padj < 0.05) between day 17 TGF-β-treated NK cells and cells from the withdrawal condition. **(B)** Heatmap showing z-score of regularized counts of TGF-β-regulated genes (defined in Fig. S3A; |log2(FC)| > 0.58, padj < 0.05) across individual donors and conditions, with k-means clustering (k = 4). Violin plots depict the same data with donors pooled by condition (n = 5). **(C)** Proportional Euler diagram showing the overlap of DARs identified by ATAC-seq between two pairwise comparisons: day 17 TGF-β vs. day 17 IL-15 (darkblue) and day 17 TGF-β withdrawal vs. day 17 IL-15 (dashed line). Each set contains significantly DA chromatin peaks (padj < 0.05, |log2(FC)| > 0.58). Numbers indicate the count of unique peaks in each area. Overlapping regions represent peaks shared between both comparisons. Peak identity is based on unique PeakIDs. **(D)** Heatmap showing z-score-normalized chromatin accessibility of TGF-β-regulated DARs (defined in Fig. S3A; |log2(FC)| > 0.58, padj < 0.05) across individual donors and conditions, with k-means clustering (k = 4). Violin plots show the same data with donors pooled by condition (n = 5).

Epigenetic profiling revealed extensive chromatin remodeling in both continuously TGF-β-treated and TGF-β-withdrawn NK cells, with 27,798 DARs detected in continuously treated NK cells and 9,182 DARs persisting after TGF-β withdrawal (**Fig. 3C).** Most DARs retained after withdrawal overlapped with regions altered under continuous exposure, indicating that withdrawal largely preserves the TGF-β-imprinted chromatin landscape (**Fig. 3C**). Consistent with transcriptional patterns, chromatin accessibility changes segregated into four temporal clusters, mirroring transient versus persistent regulation (**Fig. 3D**). Importantly, approximately 25 % of DARs detected at day 7 remained altered after TGF-β withdrawal, with the majority exhibiting sustained loss of accessibility. These persistently less accessible regions mapped to genes critical for NK cell effector function and downstream signaling, including IL12RB, GZMB, IFNG, NFKB, and IRF2 (**Fig. 3D**). Additional transcription start site (TSS)-focused analysis demonstrated sustained loss of accessibility at selected effector promoters after TGF-β withdrawal, notably GZMA, IFNG, GZMM, and PRF1, consistent with a TGF-β-driven mechanism that locks specific TSSs in a less accessible state (**Fig. S3 B-C**). Together, these findings suggest that prolonged TGF-β exposure establishes a durable epigenetic state characterized by persistent restriction of effector-associated regulatory regions that is maintained independently of continued signaling.

Given the persistence of reduced chromatin accessibility at effector loci after TGF-β withdrawal, we next asked whether strong reactivation signals could restore NK cell function by at least partially reversing the underlying epigenetic changes. To this end, we applied (i) prolonged cytokine-driven activation using cytokine combinations commonly used to generate CIML NK cells ^21, 24^, and (ii) sequential stimulation with activating receptor ligation followed by cytokines (**Fig. S3D**), based on the rationale that prior receptor engagement can epigenetically prime NK cells and reshape subsequent cytokine-driven transcriptional responses ^27^. However, neither approach resulted in recovery of effector function (**Fig. S3E**), suggesting that the TGF-β-induced state is resistant to reactivation. These findings indicate that prolonged TGF-β exposure establishes an epigenetically stabilized state that constrains re-engagement of NK cell effector programs despite strong activating signals.

### Durable epigenetic repression targets IRF-, EOMES-, and T-bet-associated regulatory elements

To elucidate the mechanisms underlying durable epigenetic repression at some loci but not others, we performed TF motif analysis across the distinct ATAC-seq accessibility clusters. Regions exhibiting sustained loss of accessibility after prolonged TGF-β exposure were strongly enriched for binding motifs of IRFs as well as the lineage-defining TFs EOMES and T-bet (**Fig. 4A**), indicating preferential repression of regulatory elements central to NK cell identity and effector function ^28, 29, 30, 31^ ^32, 33^. In contrast, regions that gained accessibility were enriched for CTCF and BORIS motifs (**Fig. S4A**), suggesting altered higher-order chromatin organization ^34, 35^. Regions that were transiently less accessible at day 7 but recovered after signal withdrawal were enriched for NF-κB family motifs (**Fig. S4A**), consistent with reversible suppression of activation-associated transcriptional programs^36^.

**Figure 4.**
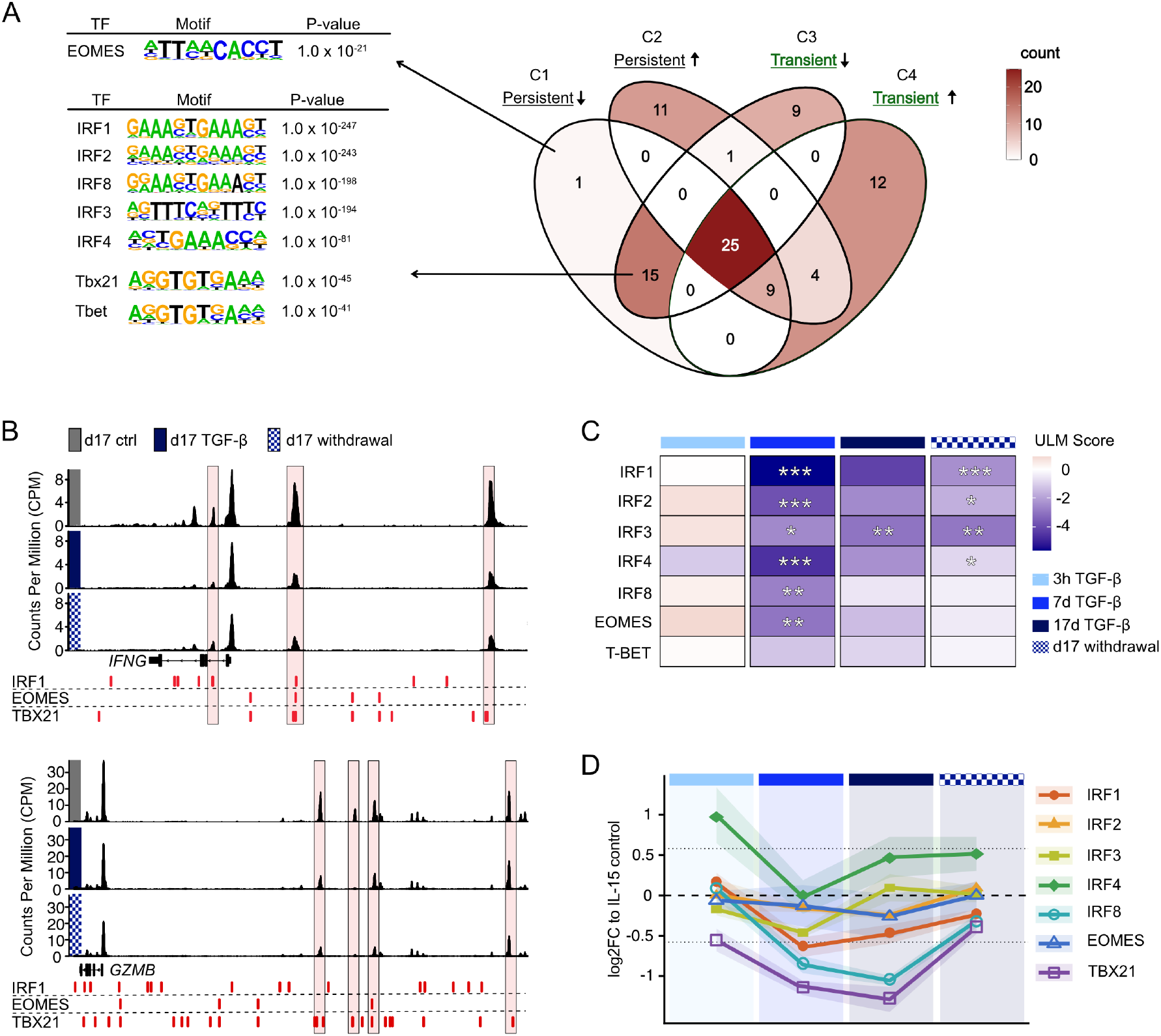
IRF and lineage-defining TF motifs are enriched in persistently closed chromatin regions. ATAC-seq was performed on primary human NK cells at day 17 following stimulation with IL-15, TGF-β, or TGF-β for 7 d followed by 10-day culture in IL-15 (TGF-β withdrawal). **(A)** TF motif enrichment analysis of ATAC-seq regions previous analysis (Fig.3D, C1, C2, C3, C4). The Venn diagram depicts the overlap of the top 50 enriched TF motifs (ranked by p value) across the indicated region categories; color intensity reflects the number of TFs assigned to each region. HOMER analysis was performed on pre-filtered regions (persistent down/up; transient down/up). For TFs present in more than one region, the most significant p value is shown. **(B)** Genome browser tracks of ATAC-seq signal at the IFNG and GZMB loci from one representative donor, shown as normalized read coverage. Predicted TF binding sites within promoter and putative enhancer regions are indicated. Boxes highlight regions of interest. **(C)** Transcription factor (TF) activity was inferred using the Univariate Linear Model (ULM) method from the decoupleR framework, applied to differentially expressed genes (DEGs) from each pairwise condition comparison. ULM scores reflect the estimated activity of each TF based on the enrichment of its target genes within the DEG set. Only TFs with positive mode-of-regulation (MOR) interactions and more than 10 target genes in the TGFβ-regulated gene set were included. Significance of TF activity is indicated as * p < 0.05, ** p < 0.01, *** p < 0.001. **(D)** Gene expression was quantified from raw counts using DESeq2 size-factor normalization. Values are shown as log2 fold change relative to the time-matched IL-15 control. Lines represent the mean ± SEM across replicates (n = 5).

To assess the functional relevance of these motif enrichments, we examined DARs at key effector gene loci. DARs linked to *IFNG, GZMA* and *GZMB* harbored binding sites for IRF1, EOMES, and T-bet, and exhibited sustained loss of accessibility following prolonged TGF-β exposure (**Fig. 4B, Fig. S4B)**. Given the high similarity of IRF family consensus binding motifs, IRF1 binding sites are shown as representative of the broader IRF family in genomic track visualizations. Because IRF family members integrate inflammatory cytokine signaling while EOMES and T-bet sustain NK cell lineage identity and effector competence, persistent restriction of these regulatory regions provides a mechanistic basis for durable impairment of NK cell activation and responsiveness ^29, 37, 38, 39^ ^31, 40, 41, 42, 43, 44, 45^.

We next inferred TF activity across treatment conditions using univariate linear models (ULM) applied to the CollecTRI regulon network ^46, 47^. Activity of multiple IRF family members (IRF1, IRF2, IRF3, IRF4, IRF8), as well as EOMES but not T-bet, was significantly reduced after 7 days of TGF-β exposure (**Fig. 4C**). Although partial recovery of several factors was observed by day 17 and after TGF-β withdrawal, none returned to baseline activity levels: IRF1, IRF2, IRF3 and IRF4 activity remained significantly suppressed in the withdrawal conditions, while IRF8 and EOMES showed a trend toward reduced activity that did not reach statistical significance, likely reflecting increased variance at later timepoints rather than genuine recovery (**Fig. 4C**).

Analysis of transcript abundance showed that reduced inferred TF activity was only partly mirrored at the mRNA level. *TBX21*, *IRF1* and *IRF8* were downregulated during prolonged TGF-β exposure but recovered after withdrawal, indicating that reduced activity after withdrawal cannot be explained by sustained transcriptional repression. In contrast, *EOMES* and *IRF2* remained largely unchanged at the transcript level, despite reduced inferred activity and enrichment of their motifs in persistently less accessible regions (**Fig. 4D**). Thus, persistent impairment of effector-associated TF programs is not primarily explained by loss of TF expression, but instead reflects altered accessibility and engagement of their regulatory targets.

Beyond these factors, we also observed persistently reduced activity of STAT1, STAT3, STAT4, and STAT5B at day 17 after signal withdrawal (**Fig. S4C**), suggesting broader disruption of cytokine-responsive transcriptional networks beyond the TFs identified by motif enrichment.

Taken together, these findings demonstrate that prolonged TGF-β exposure selectively disrupts transcriptional networks that integrate inflammatory signaling and sustain NK cell effector identity, thereby establishing a mechanistic basis for durable multi-pathway dysfunction.

### Histone modifications underlie reversible tissue adaptation but not persistent effector dysfunction

To assess histone modification dynamics at day 17 of TGF-β exposure, we integrated H3K4me3 and H3K27ac Cleavage Under Targets and Release Using Nuclease (CUT&RUN)^48^ signals with our ATAC-seq accessibility clusters (**Fig. 5A**). At persistently less accessible regions (C1), neither H3K4me3 nor H3K27ac signals were substantially altered across conditions, remaining low irrespective of TGF-β exposure or withdrawal (**Fig. 5A**), indicating that persistent chromatin closure is not accompanied by measurable changes in either permissive histone mark. In contrast, transiently less accessible regions (C3) displayed dynamic and fully reversible changes in both H3K4me3 and H3K27ac occupancy. Regions gaining accessibility - both persistently (C2) and transiently (C4) - showed corresponding gains in H3K4me3 with no substantial changes in H3K27ac occupancy (**Fig. 5A**), indicating that TGF-β-driven chromatin opening is globally more strongly associated with H3K4me3 than with H3K27ac remodeling. Notably, transiently less accessible regions exhibited discordant H3K4me3 and H3K27ac dynamics, suggesting partial uncoupling of these active histone marks at dynamically regulated loci. Together, these findings indicate that permissive histone mark remodeling at day 17 is largely restricted to dynamically accessible regions, while the persistent epigenetic changes at effector loci occur independently of H3K4me3 and H3K27ac.

**Figure 5.**
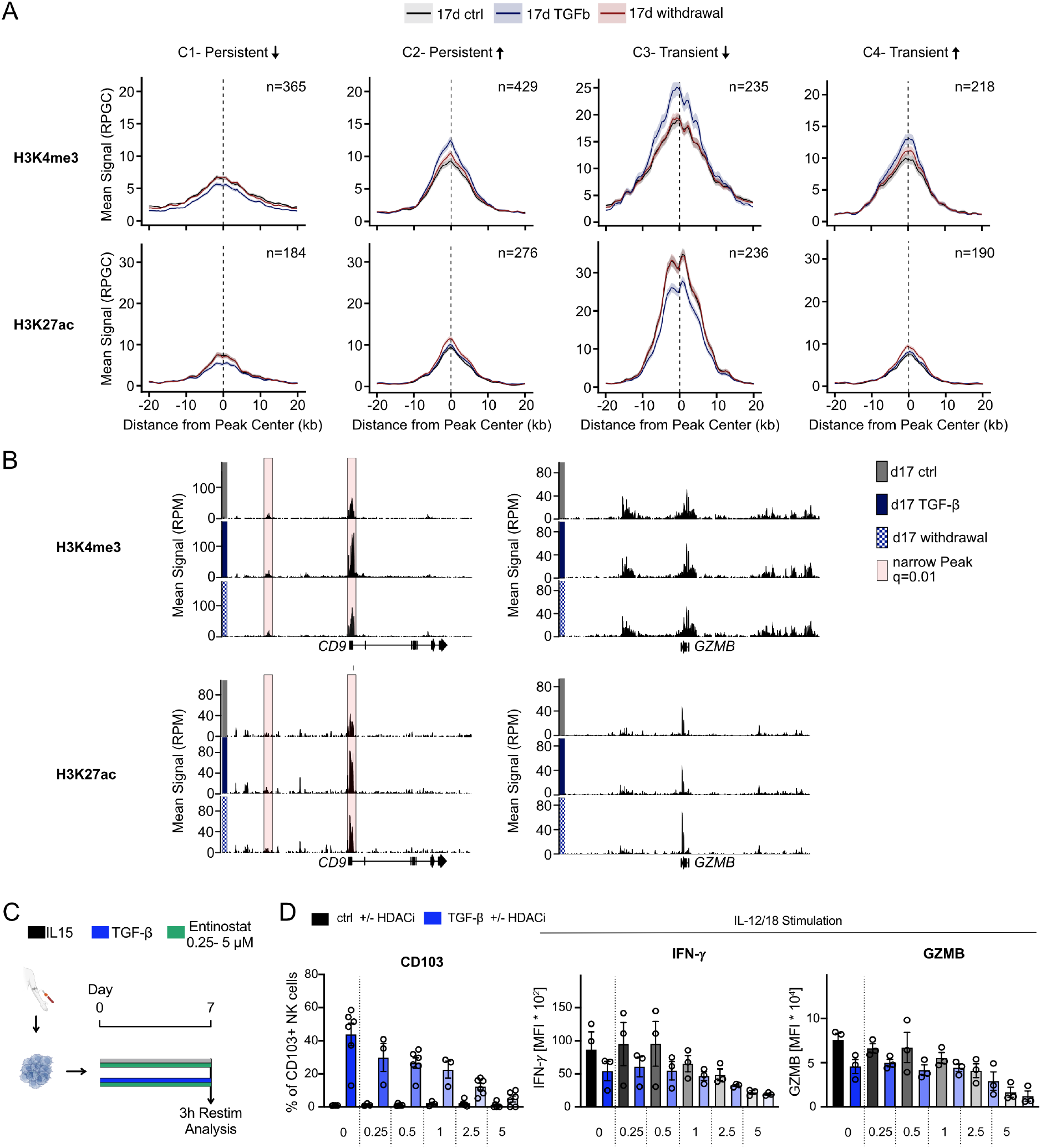
TGF-β dynamically remodels active histone marks in NK Cells. (A-B) CUT&RUN profiling of H3K4me3 and H3K27ac was performed on primary human NK cells at day 17 following stimulation with IL-15, TGF-β, or TGF-β for 7 d followed by 10-day culture in IL-15 (TGF-β withdrawal). **(A)** Mean CUT&RUN signal (RPGC) for H3K4me3 (top row) and H3K27ac (bottom row) is shown centered on peak loci across four ATAC-seq-defined chromatin clusters: C1, C2, C3, C4. Profiles represent the mean ± SEM across three biological replicates and are plotted over a ±20 kb window centered on the peak center (n=3). Anchor regions were defined using a shared peak set: consensus peaks (present in ≥ 2 of 3 replicates per condition) from all seven conditions were merged into a union set, retaining only peaks with ≥ 50% reciprocal overlap with cluster-specific ATAC-seq peaks. Signal was extracted from RPGC-normalized BigWig files in 50 bp bins. Lines represent 17 d control (gray), 17 d TGF-β (blue), and 17 d withdrawal (red). The number of anchor peaks per cluster is indicated (n). **(B)** Mean CUT&RUN signal (RPM) for H3K4me3 (top) and H3K27ac (bottom) is shown at the *CD9* (left) and *GZMB* (right) loci for day 17 control (gray), day17 TGF-β (dark blue), and day 17 withdrawal (blue hatched) conditions. Each track represents the mean signal across three biological replicates. Red shaded boxes indicate narrow peaks called at q < 0.01 (MACS3). Gene models are shown below the tracks. **(C)** Primary human NK cells were isolated and cultured for 7 d in IL-15 in the presence of TGF-β and/or the HDAC inhibitor Entinostat at concentrations ranging from 0.25 to 5 µM. Cells were re-stimulated with IL-12 and IL-18 for 3 hours at day 7 prior to analysis. **(D)** Bar graphs show the frequency of CD103+ NK cells, IFN-*γ* MFI, and Granzyme B (GZMB) MFI following 3 h IL-12/18 [IL-12: 20 ng/mL; IL18: 20 ng/mL] restimulation at day 7 (n=3-6; two independent experiments) . Cells were treated with control (black) or TGF-β (blue) in the presence of increasing concentrations of Entinostat (0, 0.25, 0.5, 1, 2.5, 5 µM). Individual data points represent biological replicates; bars show mean ± SEM.

At the locus level, TR-associated genes *CD9* and *ITGAE* displayed clear TGF-β-dependent gains in both H3K4me3 and H3K27ac that resolved after signal withdrawal, consistent with reversible transcriptional activation (**Fig. 5B**, **Fig. S5A**). In contrast, the effector loci *GZMA* and *GZMB* showed no comparable histone mark dynamics, with H3K4me3 and H3K27ac signal remaining largely unchanged across conditions, apart from one locus upstream of *GZMA* displaying transient loss in H3K27ac (**Fig. 5B, Fig. S5A**). Thus, reversible TR-associated remodeling and persistent effector restriction are associated with fundamentally distinct chromatin signatures.

To functionally test whether histone acetylation contributes differentially to TR and effector programs, we treated NK cells with increasing doses of the histone deacetylase (HDAC) inhibitor Entinostat ^49^ during TGF-β exposure (**Fig. 5C**). Entinostat dose-dependently reduced the frequency of CD9+ and CD103+ NK cells in TGF-β conditions, suggesting that TR programming is sensitive to perturbation of HDAC activity (**Fig. 5D**, **Fig. S5B**). In contrast, IFN-γ and GZMB expression were unaffected across the full dose range, indicating that effector dysfunction is not readily reversed by HDAC inhibition (**Fig. 5D**). Cell viability was preserved under Entinostat doses up to 1 µM, whereas higher Entinostat concentrations (2.5–5 μM) significantly reduced cell viability (**Fig. S5C**). Therefore, conclusions should primarily be drawn from the 0.25 - 1 μM dose range. Together, these findings reveal distinct epigenetic architectures underlying TGF-β-induced NK cell programs: whereas TR-associated adaptation is linked to dynamic and reversible histone remodeling, persistent effector dysfunction is maintained independently of active histone mark remodeling and remains resistant to HDAC inhibition.

### SMAD4 orchestrates divergent NK cell reprogramming through direct targeting of key transcriptional regulators

To identify the direct transcriptional targets of TGF-β/SMAD signaling in NK cells, we performed CUT&RUN for SMAD3 and SMAD4. High-confidence binding sites were defined as peaks identified in at least two of three replicates at q < 0.01, yielding 2,762 consensus binding sites. These were predominantly located at promoter/TSS regions (50%), with additional binding at intronic (26%) and distal intergenic regions (20%) (**Fig. 6A**). Representative loci such as *SMAD7* and *SKI* confirmed robust and specific SMAD4 occupancy at canonical TGF-β target genes (**Fig. 6B**). HOMER motif analysis of consensus peaks identified SMAD binding motifs among the top enriched sequences, confirming on-target specificity of the CUT&RUN signal (**Fig. S6A).**

**Figure 6.**
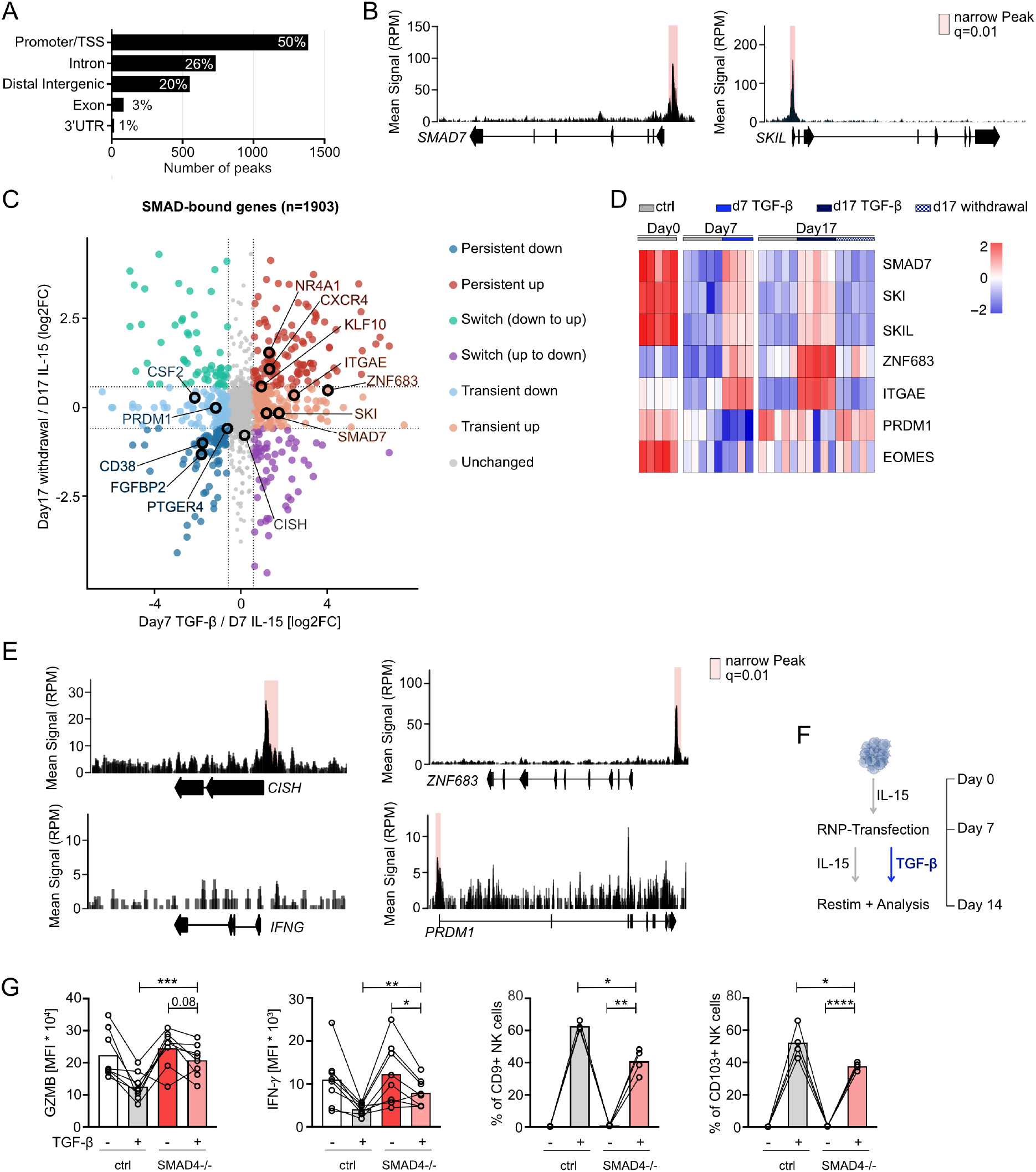
SMAD3/4 occupancy identifies divergent regulatory programs downstream of TGF-β signaling. CUT&RUN profiling of SMAD3/4 was performed on human NK cells exposed to TGF-β for 3 hours (n=3). **(A)** Genomic feature distribution of SMAD CUT&RUN peaks (n = 2,774 consensus peaks, q < 0.01) reproducible across at least two of three biological replicates (≥50% reciprocal overlap). Peaks were annotated using ChIPseeker against the hg38 genome (TSS window ± 3 kb). The majority of SMAD-binding sites localize to promoter/TSS regions (50%), followed by intronic (26%) and distal intergenic regions (20%). **(B)** Representative CUT&RUN signal tracks (mean signal in RPM) at the *SMAD7* and *SKIL* loci. Red shading indicates called SMAD-bound narrow peaks (q < 0.01; MACS3). **(C)** Scatter plot of SMAD-bound genes (n = 1,903) comparing transcriptional responses at day 7 TGF-β vs. day 7 IL-15 (x-axis, log₂FC) and at day 17 after TGF-β withdrawal vs. day 17 IL-15 (y-axis, log₂FC), derived from DESeq2 analysis of long-term TGF-β RNA-seq data. Genes are classified by response pattern using a |log₂FC| > 0.58 threshold: persistently upregulated (red), persistently downregulated (blue), transiently upregulated (salmon), transiently downregulated (light blue), switched down-to-up (teal), switched up-to-down (purple), or unchanged (grey). Selected genes of interest are labeled and highlighted with a black circle. **(D)** Heatmap of VST-normalized gene expression (variance-stabilizing transformation, DESeq2), row-scaled (z-score), for selected SMAD target genes across six conditions: unstimulated, d 7 IL-15, d 7 TGF-β, d 17 IL-15, d 17 TGF-β, and d 17 TGF-β withdrawal. Each column represents an individual biological replicate. **(E)** CUT&RUN signal tracks (mean RPM) at three SMAD target loci *CISH*, *ZNF683*, and *PRDM1*, as well as the *IFNG* locus. Red shading marks called SMAD narrow peaks (q < 0.01, MACS3). **(F)** Experimental scheme. Primary NK cells were expanded in IL-15 starting at day 0, followed by CRISPR-RNP transfection at day 7. Control cells underwent electroporation without loaded Cas9 enzyme. From day 7 to day 14, cells were cultured in IL-15 ± TGF-β, then restimulated and analyzed at d 14. **(G)** Protein expression in control (ctrl) and SMAD4-deficient (SMAD4⁻/⁻) NK cells with our without 7 d TGF-β treatment (− or +). Shown are GZMB and IFN-γ MFI, and percentage of CD9⁺ and CD103⁺ NK cells. Each connected line represents a paired biological replicate (two independent runs). Statistical significance was assessed by paired student t-test comparison; * p < 0.05, ** p < 0.01, *** p < 0.001, **** p < 0.0001.

Integration of SMAD4 binding with transcriptional dynamics revealed that SMAD-bound genes segregated into distinct persistence categories, demonstrating that direct SMAD binding does not uniformly predict the durability of transcriptional regulation (**Fig. 6C**).

A subset of SMAD-bound targets, including *ZNF683* and *ITGAE*, showed transient induction during TGF-β exposure that resolved after withdrawal, consistent with reversible regulation of TR-associated programs. In contrast, other direct targets displayed distinct temporal dynamics, including transient repression of *PRDM1* and early induction of the cytokine signaling regulator *CISH* ^50^ (**Fig. 6C–D, Fig. S6B**). These findings indicate that direct SMAD4 targets are dynamically and differentially regulated downstream of TGF-β signaling.

Notably, direct SMAD4 binding was observed at TFs such as *ZNF683, PRDM1*, and *EOMES* as well as cytokine signaling regulators including *CISH*, but was not detected at canonical effector loci such as *IFNG* and *GZMB*, despite their persistent loss of accessibility and function (**Fig. 6E**). These findings suggest that canonical TGF-β signaling influences persistent effector dysfunction indirectly through upstream regulatory programs rather than through direct SMAD4 occupancy at effector loci.

To determine whether SMAD4 is required for TGF-β-driven NK cell reprogramming, we generated SMAD4-deficient human NK cells by CRISPR/Cas9 ribonucleoprotein (RNP) electroporation and exposed them to TGF-β for seven days prior to functional analysis (**Fig. 6F**). SMAD4 deficiency rescued TGF-β-induced suppression of GZMB and partially restored IFN-γ expression, confirming that effector silencing is SMAD4-dependent (**Fig. 6G**). Conversely,

TGF-β-driven induction of CD9 and CD103 was partially reduced in SMAD4-deficient cells, indicating that TR-associated programming is at least partly dependent on intact SMAD4 signaling (**Fig. 6G).** Together, these findings suggest that SMAD4 functions as a key upstream regulator of TGF-β-driven NK cell reprogramming, promoting reversible tissue adaptation programs while indirectly shaping persistent effector dysfunction through upstream regulatory network remodeling.

### NK cells from patients with liver tumors recapitulate epigenetic and functional hallmarks of prolonged TGF-β signaling

One physiological context in which NK cells are likely to experience prolonged TGF-β exposure is the tumor microenvironment^51^. To assess whether the chromatin and transcriptional programs defined under sustained TGF-β exposure are reflected in human disease, we analyzed NK cell phenotype, function and chromatin accessibility in patients with primary liver cancer or liver metastases who were not undergoing immunotherapy (**Fig. S7A**). Tumor-infiltrating NK cells from these patients displayed significantly increased expression of CD103 (**Fig. 7A**), consistent with ongoing TGF-β signaling within the tumor. In parallel, peripheral blood NK cells from HCC patients showed reduced IFN-γ production and diminished degranulation upon restimulation, as measured by CD107a expression (**Fig. 7B**), indicating sustained functional suppression beyond the tumor site.

**Figure 7.**
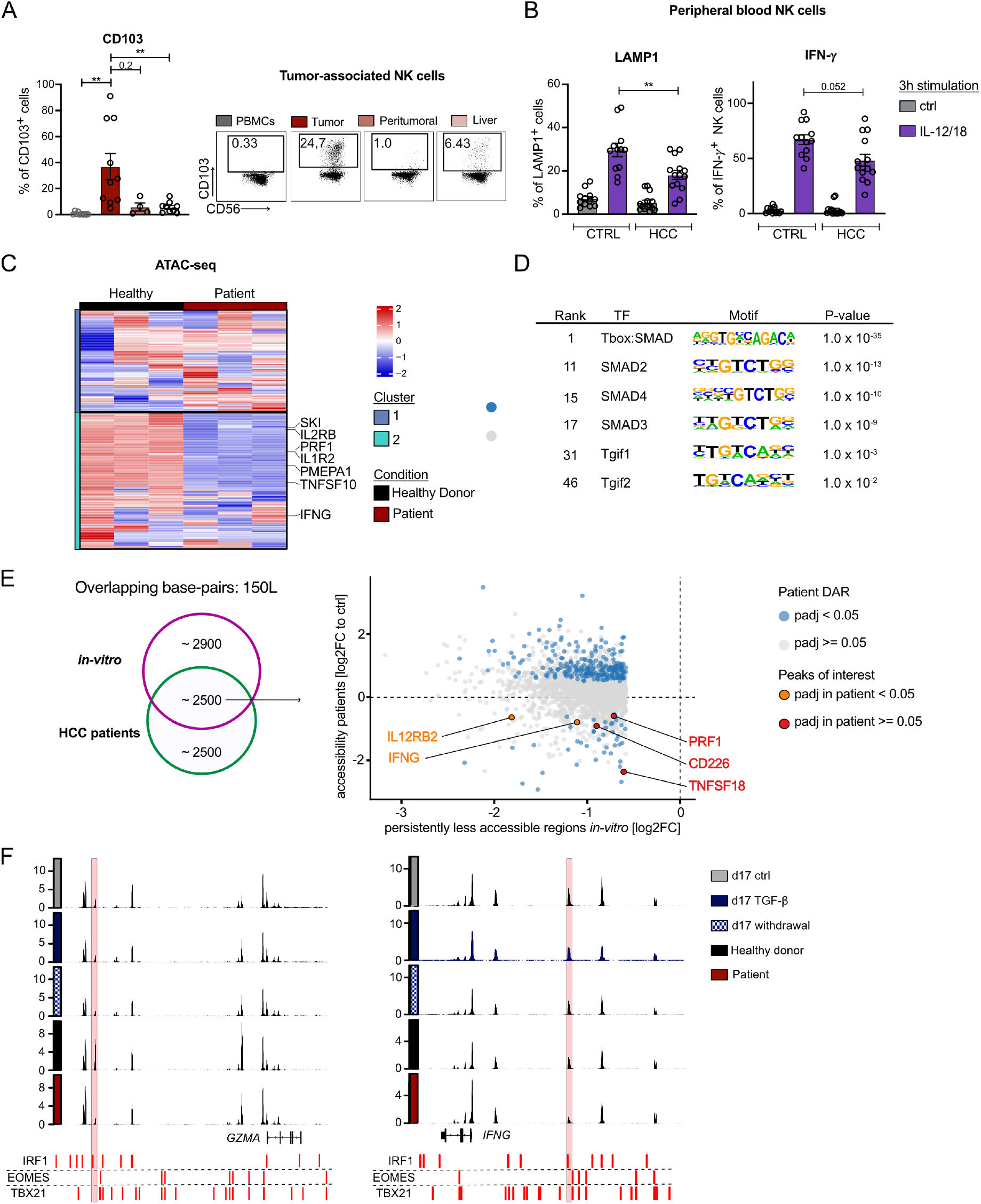
NK cells from patients with liver tumors recapitulate functional and epigenetic features of prolonged TGF-β exposure. Intratumoral NK cells from liver resections and peripheral NK cells from HCC patients were analyzed by flow cytometry and ATAC-seq. **(A)** Frequency of CD103⁺ cells among NK cells isolated from PBMCs, tumor, peritumoral tissue, and liver of patients with liver cancer or liver metastasis, shown as percentage of CD103⁺ cells. Representative flow cytometry plots showing CD103 vs. CD56 expression in the indicated compartments, with percentages of CD103⁺CD56⁺ NK cells indicated (right). **p < 0.01 (Wilcoxon test). **(B)** Peripheral NK cells from HCC patients were cultivated overnight (10ng/mL IL15; 25 U/mL) and restimulated for 4 hours with IL-12/18 (50 ng/mL both). Degranulation (LAMP1, left) and cytokine production (IFN-γ, right) are shown as bar plot. Each dot represents one donor/patient. **p < 0.01 (Wilcoxon test). **(C)** ATAC-seq was performed on sorted peripheral blood NK cells of three HCC patients with dysfunctional NK cells. Heatmap of chromatin accessibility across the top 500 most variable ATAC-seq peaks (by row variance across VST-normalized counts) comparing unstimulated NK cells from healthy donors and HCC patients (n=3 each). Selected genes of interest annotated to the respective peaks are indicated. **(D)** Transcription factor motif enrichment analysis of chromatin regions less accessible in HCC patients compared to healthy donors. Selected motifs are shown with their rank, motif logo, and associated p-value. **(E)** Scatter plot showing the concordance of chromatin accessibility changes between persistently less accessible regions identified in vitro (x-axis, log2FC of 17 d TGF-β-treated vs. day 17 control NK cells, restricted to the persistent-down cluster) and DARs in HCC patients (y-axis, log2FC of unstimulated patient vs. control NK cells). Overlapping peaks were identified by genomic coordinate overlap (minimum 150 bp). Each dot represents one overlapping peak pair, colored by significance in the patient dataset (blue: padj < 0.05, grey: padj ≥ 0.05). Selected genes of interest are highlighted and labeled; peaks labeled in red (padj < 0.05 in patients) and orange (padj ≥ 0.05). **(F)** Genome browser tracks showing chromatin accessibility (ATAC-seq signal) at the *GZMA* and *IFNG* loci across indicated conditions. Predicted binding sites for IRF1, EOMES, and T-bet transcription factors are indicated below each track in red. A binding site was called when the PWM match score reached ≥90% of the theoretical maximum score for that motif.

To determine whether this functional impairment was accompanied by stable epigenetic changes, we performed ATAC-seq on peripheral blood NK cells from patients with HCC and healthy controls. Chromatin accessibility profiles clearly distinguished patient-derived NK cells from healthy controls (**Fig. S7B)** and revealed a widespread loss of chromatin accessibility in HCC-associated NK cells (**Fig. 7C**), supporting the presence of disease-associated epigenetic repression. Consistent with impaired effector function, regions with reduced accessibility included loci associated with NK cell activation, cytotoxicity and cytokine production, including *IFNG*, *PRF1*, *IL2RB*, and *TNFSF10*. In addition, reduced accessibility was observed at SMAD target genes identified *in vitro*, including *SKI*, *IL1R2*, and *PMEPA1*, indicating that TGF-β-responsive regulatory elements are similarly altered in patient-derived NK cells.

Motif enrichment analysis of regions with decreased accessibility in patient NK cells revealed overrepresentation of SMAD and TGIF binding motifs (**Fig. 7D**), consistent with engagement of TGF-β-associated transcriptional programs *in vivo*. Notably, patient-derived DARs overlapped with regions that remained persistently less accessible following prolonged TGF-β exposure *in vitro* (**Fig. 7E**), including regulatory regions associated with *PRF1* and *IFNG*. Inspection of selected loci linked to NK cell effector function further revealed reduced chromatin accessibility at upstream regulatory regions of *IFNG*, and *GZMA* in patient-derived NK cells. These regions contained putative T-bet and IRF1 binding motifs and overlapped with TGF-β-associated DARs identified in our *in vitro* model (**Fig. 7E**), supporting the relevance of this epigenetic program *in vivo*. Additional accessibility changes not observed *in vitro* were also detected, consistent with the presence of multiple suppressive signals within the tumor environment.

Collectively, these findings demonstrate that NK cells from HCC patients acquire a disease-associated epigenetic state that recapitulates key functional and chromatin features of prolonged TGF-β exposure in vitro, supporting chronic TGF-β signaling as a central driver of durable NK cell dysfunction in human cancer.

## DISCUSSION

TGF-β is a major suppressive signal in tissues and tumor microenvironments ^52^ and TGF-β blockade is actively pursued as a therapeutic strategy in cancer, based on the premise that suppressive signaling restrains immune cell function in a reversible manner ^53^. Yet with regard to NK cells, most studies have focused on the acute effects of TGF-β, leaving unresolved whether established NK cell dysfunction depends on continued signaling or persists after signal withdrawal ^12, 13, 14, 18, 19^. This distinction is critical: if chronic TGF-β exposure imprints a stable dysfunctional state, late therapeutic intervention may fail to restore NK cell activity. Here, we demonstrate that prolonged TGF-β signaling enforces durable epigenetic remodeling and sustained effector suppression, whereas short-term exposure induces largely reversible changes, revealing that the reversibility of NK cell dysfunction is fundamentally determined by the duration of signaling. Importantly, the chronically imprinted state induced by prolonged TGF-β signaling is refractory to diverse reactivation cues, including proinflammatory cytokines and activating receptor engagement and combinations of these, indicating a loss of functional plasticity rather than simple signal-dependent suppression.

A central mechanistic insight of our study is that prolonged TGF-β exposure selectively remodels distinct NK cell regulatory programs rather than uniformly reshaping the epigenome. While the majority of TGF-β-responsive genes, including TR-associated genes, recover chromatin accessibility and transcription upon signal withdrawal, a substantial subset of loci remains epigenetically altered. Notably, regions that fail to regain accessibility are highly enriched for binding motifs of IRFs as well as the lineage-defining TFs EOMES and T-bet, which together integrate cytokine signaling and maintain NK cell effector identity^28, 29, 31, 37, 40, 42, 44, 45^. Persistent closure of IRF-, EOMES-, and T-bet-associated regulatory elements provides a mechanistic basis for durable effector repression and loss of responsiveness to activating stimuli.

Importantly, persistent dysfunction was not accompanied by sustained changes in canonical permissive histone marks. Although H3K4me3 and H3K27ac dynamically responded to TGF-β exposure, both largely reverted after withdrawal, even at loci that remained functionally impaired. Consistent with this, HDAC inhibition preferentially affected TR-associated programs but failed to restore effector function, suggesting that durable dysfunction is maintained primarily through stable chromatin accessibility remodeling rather than persistent changes in permissive histone modifications.

Our data further indicate that TGF-β canonical signaling does not directly repress effector loci, but instead rewires upstream transcriptional circuitry. SMAD3/4 CUT&RUN identified direct binding at transcriptional regulators and cytokine response genes including *ZNF683*, *PRDM1*, and *CISH*, whereas canonical effector loci such as *IFNG* and *GZMB* lacked detectable SMAD occupancy. Notably, the reciprocal and transient regulation of *ZNF683* (Hobit) and *PRDM*1 (Blimp-1) is consistent with control of a reversible tissue adaptation module downstream of canonical TGF-β signaling^54^. Together with the partial rescue observed in SMAD4-deficient NK cells, these findings support a model in which canonical TGF-β signaling bifurcates NK cell fate into two mechanistically distinct programs: a dynamically reversible tissue adaptation program and a more durable suppression of effector competence associated with stable epigenetic remodeling.

Our findings extend previous work proposing largely SMAD4-independent regulation of TR programs^55^ by demonstrating that canonical SMAD4 signaling directly contributes to TR-associated transcriptional regulation. These results appear at odds with findings in mice, where SMAD4 restrains a non-canonical TGF-β-dependent pathway, such that SMAD4 deficiency enhances TGF-β-driven ILC1-like conversion in NK cells^26^. Rather than reflecting a fundamental species difference, these divergent findings may instead point to context-dependent engagement of canonical versus non-canonical TGF-β circuits. In line with this model, SMAD4 occupied upstream TR regulators including *ZNF683* and *PRDM1* in our system, whereas loss of SMAD4 only partially reduced CD103 and CD9 expression, indicating that non-canonical pathways likely contribute in parallel.

More broadly, our findings support the concept that the functional consequences of TGF-β signaling are highly context-dependent. Chronic TGF-β exposure during strong concurrent activation has been proposed to promote cytokine hypersecretion rather than suppression^56^, indicating that activating cues can fundamentally reshape downstream TGF-β responses.

Similarly, Horowitz and colleagues showed that transient tumor cell interactions drive differentiation into cytotoxic tissue-resident NK cells in a TGF-β-dependent manner, whereas sustained exposure to soluble TGF-β alone induces a hyporesponsive CD9+ decidual-like state closely resembling the dysfunctional program identified here^57^. These observations converge with our own to suggest that signal duration and co-stimulatory context critically shape divergent TGF-β-driven NK cell fates, spanning reversible tissue adaptation, cytotoxic tissue residency, and epigenetically stabilized dysfunction.

Our findings further place durable NK cell dysfunction within the wider framework of chronic lymphocyte dysfunction in persistent infection and cancer. In T cells, prolonged TGF-β signaling contributes to exhausted or attenuated states associated with reduced T-bet and EOMES activity ^58, 59, 60^ ^61, 62^, yet these programs often remain at least partly reversible following pathway blockade ^58, 59, 60, 63, 64^. In contrast, our data indicate that prolonged TGF-β exposure can impose a more durable state in NK cells that persists independently of continued signaling. Although recent work in AML implicated a TGF-β-SMAD-BATF axis in prolonged NK cell dysfunction^20^, BATF was neither induced nor directly targeted by SMAD4 in our system. Instead, our data identify stable restriction of IRF-, EOMES-, and T-bet-associated regulatory elements as a defining feature of chronically imprinted NK cell dysfunction, linking prolonged TGF-β exposure to durable impairment of core NK cell effector programs.

Our observations further suggest that prolonged exposure to suppressive tissue environments may leave durable epigenetic imprints on circulating NK cells even after tissue egress. Tissue-resident NK cells can re-enter circulation and exhibit phenotypic plasticity ^65^. Consistent with this model, peripheral blood NK cells from HCC patients exhibit impaired effector function together with marked epigenetic remodeling overlapping with features induced by prolonged TGF-β exposure *in vitro*. Similar dysfunctional CD9+ NK cell states have recently been described in PDAC, where chronic TGF-β-rich tumor environments promote decidual-like NK cell polarization associated with impaired NK cell function, and poor clinical outcome^66^.

Together, these observations support the concept that chronic TGF-β exposure in human tumors can durably imprint dysfunctional NK cell states *in vivo*.

These insights have important therapeutic implications. Our data suggest that once prolonged TGF-β exposure has established stable epigenetic remodeling at effector-associated regulatory elements, removal of the suppressive signal alone is insufficient to restore NK cell competence. This may help explain the limited efficacy of late TGF-β pathway inhibition or cytokine restimulation strategies. Consistent with this, pharmacologic inhibition of TGF-β signaling with galunisertib^67^ has shown limited efficacy in restoring NK cell activity as a standalone approach and may require combination with adoptive NK cell transfer^20^. Accordingly, our findings suggest that successful NK cell reinvigoration may require therapeutic strategies that target both suppressive signaling pathways and the underlying imprinted chromatin state.

Several limitations should be acknowledged. Our study relied on controlled *in vitro* systems that cannot fully recapitulate the complexity of tissue microenvironments *in vivo*, and prolonged IL-15 exposure may itself contribute to transcriptional and epigenetic adaptation, although analyses were restricted to TGF-β-associated changes wherever possible. In addition, although our data identify stable loss of accessibility at IRF-, EOMES-, and T-bet-associated regulatory regions as a hallmark of persistent dysfunction, the molecular mechanisms that maintain these inaccessible states remain to be resolved in detail.

Collectively, our findings identify TGF-β signal duration as a critical determinant of NK cell fate that separates reversible tissue adaptation from epigenetically stabilized effector dysfunction. More broadly, these data support a model in which chronic suppressive signaling can durably encode dysfunctional lymphocyte states through selective epigenetic remodeling, with important implications for the timing and design of NK cell-directed immunotherapies.

## MATERIALS AND METHODS

### Study design

This study investigates how the duration of TGF-β exposure shapes NK cell transcriptional, epigenetic, and functional states. Primary human NK cells were exposed to short-term or prolonged TGF-β stimulation followed by withdrawal and restimulation conditions to assess reversibility of induced programs. Functional assays, RNA-seq, ATAC-seq, CUT&RUN, and CRISPR-RNP-mediated knockout were used to define associated molecular changes, and patient-derived NK cells from individuals with HCC were analyzed for comparison with *in vitro*-induced dysfunctional states. Independent biological donors were used throughout as indicated in the figure legends.

### Human samples

Peripheral blood mononuclear cells (PBMCs) were collected from healthy adult volunteers. The study was reviewed and approved by the Ethics Committee of the Technical University of Munich (TUM), Germany (approval number 2025-584-S-NP). PBMCs were isolated using a Pancoll density gradient. NK cells were enriched using an NK Cell Isolation Kit (Miltenyi). Cells were cultured in complete IMDM GlutaMAX (Gibco) medium supplemented with penicillin (100 U/mL) and streptomycin (100 µg/mL) (Thermo Fisher), 1× non-essential amino acids (Thermo Fisher), and 10% FCS (Thermo Fisher), and further supplemented with recombinant human IL-15 (50 ng/mL; Miltenyi).

### Cultivation of NK cells

Depending on cell numbers, NK cells were distributed across 6-12 wells of a 96-well plate. NK cells were cultivated in different conditions as described in the text. In general, they were either cultured with IL-15 (50 ng/mL, Miltenyi) alone or in combination with IL-15 and TGF-β (10 ng/mL, R&D) in IMDM+ at 37°C, 5% CO2. In cases where TGF-β was withdrawn, cells were transferred to a low-bind tube, washed once with PBS, resuspended in fresh media, and transferred to a new well. All procedures were performed in U-bottom plates. For the 17-day setup, cells were neither handled nor supplemented between days 0-7 and 7-17.

### Immunophenotyping by multiparametric flow cytometry

At each experimental endpoint, NK cells were washed with PBS (Thermo Fisher) and stained for 30 min at 4 °C with directly labeled antibodies against extracellular markers. For intracellular staining, cells were fixed and permeabilized (4% PFA in PBS; Santa Cruz Biotechnology; eBioscience Permeabilization Buffer, Thermo Fisher) and then stained for intracellular antigens for 30 min at 4 °C.

For MitoTracker Green and TMRE staining, NK cells were incubated with the dyes for 15 min at 37 °C. Cells were then stained with a live/dead marker and immediately acquired by flow cytometry (Cytek Aurora).

### Restimulation and Degranulation Assay

For restimulation, NK cells were equally distributed across the different conditions. For cytokine activation, NK cells were stimulated for 3 hours with IL-12 (Miltenyi, 20 ng/mL), IL-18 (Miltenyi, 20 ng/mL), IL-2 (Miltenyi, 25 U/mL), IL-15 (Miltenyi, 50 ng/mL), or IFN-α (BioLegend, 25 ng/mL) in a U-bottom plate.

For plate-bound stimulation, a 96-well flat-bottom plate was coated with anti-NKp46 and/or anti-NKG2D antibodies (Miltenyi, both 10 µg/mL) for 1 hour, washed three times with PBS, and NK cells were seeded. Stimulation was initiated by a brief centrifugation for 5 seconds at 100x g.

After 1 hour of stimulation, anti-LAMP-1 (1:100, BioLegend) was added to the cells. Following stimulation, cells were transferred to a V-bottom plate for staining and processed as described above.

For sequential stimulation, a flat-bottom plate was first coated with anti-NKp46 and/or anti-NKG2D antibodies (Miltenyi, both at 10 µg/mL) for 1 hour, then washed three times with PBS. NK cells were then added, pelleted at 100x g for 5 seconds and stimulated for 3 hours, after which cells were transferred to a U-bottom plate, washed once, and resuspended in 100 µL of supplemented IMDM+. Subsequently, 2× cytokine mix was added and cells were incubated overnight. Following overnight stimulation, cells were transferred to new wells and resuspended in IL-15-containing media only. After 3 days, cells were again stimulated with cytokines for 3 hours and prepared for flow cytometry. As a control for the plate-bound approach, NK cells were seeded in uncoated wells.

### Bulk RNA sequencing and data processing

At the end of each time point, 50,000 NK cells were collected and sorted on a FACSAria III cell sorter (BD Biosciences). Singlets were gated, and viable CD56⁺ cells were selected using a live/dead discrimination dye. Sorted cells were subjected to RNA isolation using the Arcturus PicoPure RNA Isolation Kit (Applied Biosystems), including a DNase digestion. RNA concentration and integrity were assessed using Pico kits on a Bioanalyzer system (Agilent Biosciences). Only samples with an RNA integrity number (RIN) ≥ 8 were submitted for sequencing. Five biological replicates per condition were sequenced. RNA sequencing was performed by a commercial service provider.

Raw paired-end RNA-sequencing data (FASTQ) were processed in R (2023.09.0+463). Read quality control and trimming were performed using fastp via the Rfastp package (v 1.14.0). Reads were aligned to the human reference genome (hg38, UCSC) using the splice-aware subjunc aligner implemented in Rsubread (v 2.24.0)^68^.

Gene annotation was obtained from TxDb.Hsapiens.UCSC.hg38.knownGene (v 3.22.0) and used to generate a Simplified Annotation Format (SAF) file for annotation-guided alignment. BAM files were sorted and indexed using Rsamtools (v 2.26.0) (https://doi.org/10.18129/B9.bioc.Rsamtools). Gene-level read counts were generated with

GenomicAlignments (v 1.46.0)^69^, ignoring strand specificity. Entrez gene identifiers were mapped to gene symbols using org.Hs.eg.db, and genes without annotated symbols were excluded.

### Bulk ATAC sequencing and data processing

ATAC-seq was performed as described previously^70^. For further computational analysis the raw sequencing files were trimmed for adapter contamination and low quality bases with Trimmomatic 0.39^71^ (arguments: ILLUMINACLIP:adapters/NexteraPE-PE.fa:2:30:10:2:True LEADING:3 TRAILING:3 SLIDINGWINDOW:4:15 MINLEN:36) and aligned against the *Homo sapiens* genome assembly GRCh38.p14 using Bowtie2^72^ 2.5.0 (arguments: --end-to-end -- no-mixed --dovetail --very-sensitive -X 615). Post-alignment, mitochondrial and low-quality fragments (equivalent SAMtools view arguments: -F 2828 -f 3 -q 30) were removed using pysam 0.22.0. Fragment duplicates were filtered out using Picard 3.1.1 and alignments to regions defined in the ENCODE blacklist^73^ were removed. Broad and narrow peaks were called for each sample separately using MACS 3.0.1^74^ (arguments: -f BAMPE -q 0.05 --keep-dup all --cutoff-analysis -g 2884242992; additional broad peak arguments: --broad --broad-cutoff 0.1). All identified overlapping peaks were subsequently merged with BEDtools^75^ 2.30.0 to form a set of non-overlapping fragment pile-up regions that were annotated based on their nearest transcription start site with HOMER 5.1 before counting sample-specific fragment occurrences per individual region using featureCounts^76^ 2.0.6 (arguments: -O -p --countReadPairs). Quality control metrics were assessed using FastQC 0.12.1, Qualimap 2.3 and deepTools 3.5.6^77, 78^.

### Histone and transcription factor CUT&RUN and data processing

For CUT&RUN, between 250,000 - 500,000 sorted NK cells were used per reaction. Cells were washed with PBS and resuspended in Antibody Buffer (1X eBioscience Perm/Wash Buffer, 1X Roche cOmplete EDTA-free Protease Inhibitor, 0.5 µM Spermidine, plus 2 µM EDTA in H₂O), then incubated with the desired antibody (H3K4me3: #07-473; Merck Millipore; H3K27ac: #13-0059; EpiCypher; CTCF: #2899S; CellSignaling ;all were used at a 1:50 dilution) in 96-well V-bottom plates. After antibody binding, cells were washed twice with Buffer 1 (1X eBioscience Perm/Wash Buffer, 1X Roche cOmplete EDTA-free Protease Inhibitor, 0.5 µM Spermidine) and resuspended in 50 µL of Buffer 1 containing pA/G-MNase (Cell Signaling, cat. 57813). The mixture was incubated on ice for 1 hour, followed by two washes with Buffer 2 (0.05% w/v Saponin, 1X Roche cOmplete EDTA-free Protease Inhibitor, 0.5 µM Spermidine in PBS), repeated three times. To activate the MNase digestion, cells were resuspended in Calcium Buffer (Buffer 2 + 2 µM CaCl₂) and incubated on ice for 30 minutes. Digestion was stopped by adding an equal volume of 2X STOP Buffer (Buffer 2 + 20 µM EDTA + 4 µM EGTA) along with 1 pg of Saccharomyces cerevisiae spike-in DNA (Cell Signaling, cat. 29987). Samples were incubated at 37°C for 15 min, after which DNA was extracted and purified using the Qiagen MinElute Kit.

Data processing was done using a custom R pipeline. Paired-end sequencing reads were aligned to the human reference genome (hg38) using Bowtie2 (--dovetail, --very-sensitive, --no-mixed, -- no-discordant). Aligned reads were filtered for mapping quality (MAPQ ≥ 30), mitochondrial reads were removed, and ENCODE blacklist regions were excluded. PCR duplicates were marked using samtools fixmate and markdup. Peak calling was performed using MACS3 (-f BAMPE, -q 0.01, --keep-dup all, --SPMR, --nomodel) for both narrow and broad peak detection. BigWig coverage tracks were generated using deepTools bamCoverage (50 bp bins, RPGC normalization, effective genome size: 2,913,022,398). Narrow peaks were filtered using overlap-based filtering, retaining only peaks where at least 50% of the shorter fragment overlapped with the intersecting peak.

### CRISPR RNP

Primary NK cells were expanded for seven days in IMDM supplemented with 50 ng/mL recombinant human IL-15 prior to electroporation. On day 7, ribonucleoprotein (RNP) complexes were freshly assembled by combining 2.5 µL of Cas9-NLS protein (61 µM; IDT) with 2.5 µL of sgRNA (120 µM) and incubating for 15 min at 37°C. For targeting of SMAD4, a pre-designed cocktail of three sgRNAs was obtained from EditCo Bio, reconstituted to 120 µM, and used at the standard volume of 2.5 µL per RNP assembly reaction without adjustment of total guide volume.

For electroporation, 1 × 10⁶ NK cells per condition were pelleted by centrifugation at 300 × g for 5 min, washed once with PBS, and resuspended in 10 µL of freshly prepared P3 Nucleofector solution (Lonza; Nucleofector Solution:Supplement ratio of 4.5:1). The cell suspension was transferred to the assembled RNP mixture, followed by addition of 2 µL of a 1:10-diluted electroporation enhancer oligonucleotide (IDT). A total of 25 µL per sample was loaded into a 16-well Nucleocuvette strip, and electroporated using the Lonza 4D-Nucleofector system with program EN-138.

Immediately following electroporation, 80 µL of warm IMDM supplemented with 40–50% FCS was added per well, and cells were allowed to recover for 10 min at 37°C. Cells were subsequently transferred to 48-well plates containing 500 µL IMDM with 50 ng/mL IL-15 and cultured for 2–3 days to permit recovery. From day 7 to day 14, cells were maintained in IL-15 (50 ng/mL) in the presence or absence of TGF-β (10 ng/mL). Analyses were performed following restimulation on day 14.

### Visualization of Sequencing data

PCA analysis and differential expression or chromatin accessibility analysis were performed using DESeq2 [1.50.2]^79^. Venn diagrams were generated with eulerr [7.0.4] (https://doi.org/10.32614/CRAN.package.eulerr) or VennDiagram [1.7.3] (https://doi.org/10.32614/CRAN.package.VennDiagram), and heatmaps were created using pheatmap [1.0.13] (https://doi.org/10.32614/CRAN.package.pheatmap). Motif discovery was carried out using HOMER [5.1], while gene set enrichment analysis (GSEA) was performed using msigdbr [25.1.1] (https://doi.org/10.32614/CRAN.package.msigdbr) and visualized with clusterProfiler [version]. Transcription factor (TF) activity was estimated using the univariate linear model (ULM) implemented in decoupleR [2.16.0]^47^, based on the CollecTRI TF-target network^46^. ATAC-seq and CUT&RUN genomic tracks were visualized by importing BAM files into IGV (v2.19.5). General data manipulation was conducted using the tidyverse [2.0.0] (https://doi.org/10.32614/CRAN.package.tidyverse), with additional visualizations produced using either built-in plotting functions or ggplot2 [4.0.1] (https://doi.org/10.32614/CRAN.package.ggplot2).

### Patient samples

HCC tumor samples were obtained from patients undergoing surgical resection. (Ethics approval number 232/19 S) Tumor tissue was minced into small pieces. Tissue was digested in RPMI GlutaMAX (Thermo Fisher) supplemented with 1 mg/mL collagenase IV (Roche), 100 µg/mL DNase I (Roche), 10% FCS, and 1% Penicillin-Streptomycin at 37°C for 30 minutes.

Lymphocytes were enriched using a 40/60% Percoll gradient and centrifuged at 800x g for 20 minutes at 20°C. The interphase containing lymphocytes was collected, washed with PBS, and stained as described above for flow cytometry acquisition.

Blood from HCC patients was obtained from the HCC outpatient clinic (ethics approval numbers 232/19 S and 2025-584-S-NP). Blood NK cells were isolated as described above and cultured overnight in IMDM+ medium supplemented with 10 ng/mL hIL-15 (Miltenyi) and 25 U/mL IL-2 (Miltenyi). Cells were restimulated with IL12 [50ng/mL] (Miltenyi) and IL18 [50ng/mL] (Miltenyi) for 4 hours. Anti-LAMP1 antibody to access degranulation was added from beginning followed by Brefeldin A (1X, ThermoFisher) and Monensin (1X, ThermoFisher) after 1 hour of stimulation. Intracellular staining was done o/n and flow cytometry analysis were performed as described above.

All selected HCC patients had active disease, no current immunotherapy and no local therapy within the last 2 months. Patients selected for ATAC-seq were not on tyrosine kinase inhibitor therapy or any other systemic therapy and additionally met the above criteria.

### Statistical analyses

All data are displayed as mean ± SEM unless otherwise indicated, with each dot representing one individual human donor. For pairwise comparisons, an unpaired two-tailed Student’s t-test was used for Figures 2E, 6G, and Supplementary Figure 2F, and a Wilcoxon signed-rank test was used for all other comparisons. Statistical significance is denoted as *p<0.05, **p<0.01, ***p<0.001, and ****p<0.0001. All statistical analyses were performed using GraphPad Prism and R software.

For ATAC-seq data, peaks were called using MACS 3.0.1 with the following arguments: -f BAMPE -q 0.05 --keep-dup all --cutoff-analysis -g 2884242992. Broad peaks were additionally called with --broad --broad-cutoff 0.1. For CUT&RUN data, peak calling was performed using MACS3 with the arguments -f BAMPE -q 0.01 --keep-dup all --SPMR --nomodel. Differentially accessible regions (DARs) and differentially expressed genes (DEGs) were defined as log₂ fold change ≥ 0.58 with an adjusted p-value (padj) < 0.05.

For RNA-seq data, differential expression analysis was performed using DESeq2, applying its default Wald test for significance testing. Read counts were modeled using a negative binomial distribution, and p-values were adjusted for multiple testing using the Benjamini-Hochberg procedure. All bioinformatic analyses were performed in the R statistical computing environment.

## Supplementary Materials

Supplementary figures 1-7

## Acknowledgments

We thank all Wiedemann lab members for helpful discussions and support. We also thank Colleen Lau and Hyunu Kim for their assistance with regard to bioinformatic questions. We thank all patients and healthy donors who contributed to this work. We also thank Sam Sheppard for critical lecture of the manuscript. The authors acknowledge the use of AI-based tools to assist with language editing and improvement of manuscript clarity. The content and interpretations are the authors’ own.

## Funding

German Research Foundation (DFG) Emmy Noether grant WI-4927_2-1

(GMW) Else Kröner Fresenius Foundation (EKFS) grant 2022_EKEA.85 (GMW)

TUM Junior Fellows Fund (GMW)

German research foundation (DFG) SPP-2306 – 461704785 (JPB)

German research foundation (DFG) SFB-1479 – 441891347-P21 (JPB)

German research foundation (DFG) 424926990 (JPB)

German research foundation (DFG) 442405234 (JPB)

German research foundation (DFG) 449174900 (JPB)

Wilhelm Sander-Stiftung project number 2024.133.1 (JPB)

European Research Council (ERC) under the European Union’s Horizon Europe Framework Programme for Research and Innovation grant agreement No. 101231434 — EICO-CODE (JPB)

German Cancer Aid (Deutsche Krebshilfe) Mildred-Scheel postdoctoral fellowship (EM)

## Author contributions

Conceptualization: KS, GMW

Methodology: KS, CH, JS, EM, RS

Formal analysis: KS, RS

Resources: UB, UE, ML, NH, DH, RMS

Investigation: KS, CH, JS, EM, LM, RS, KP, JB, KB

Visualization: KS, CH, JS, EM

Funding acquisition: GMW, JPB

Supervision: GMW

Writing – original draft: GMW, KS

Writing – review & editing: GMW, KS, JS, EM, RS, UB, UE, ML, NH, DH, JPB, RMS.

## Data and materials availability

All sequencing data and code will be made publicly available upon revision.

