## Supplementary Figures for "A temporal threshold in TGF-β signaling separates reversible tissue adaptation from persistent NK cell effector dysfunction"

Supplementary Figure 1

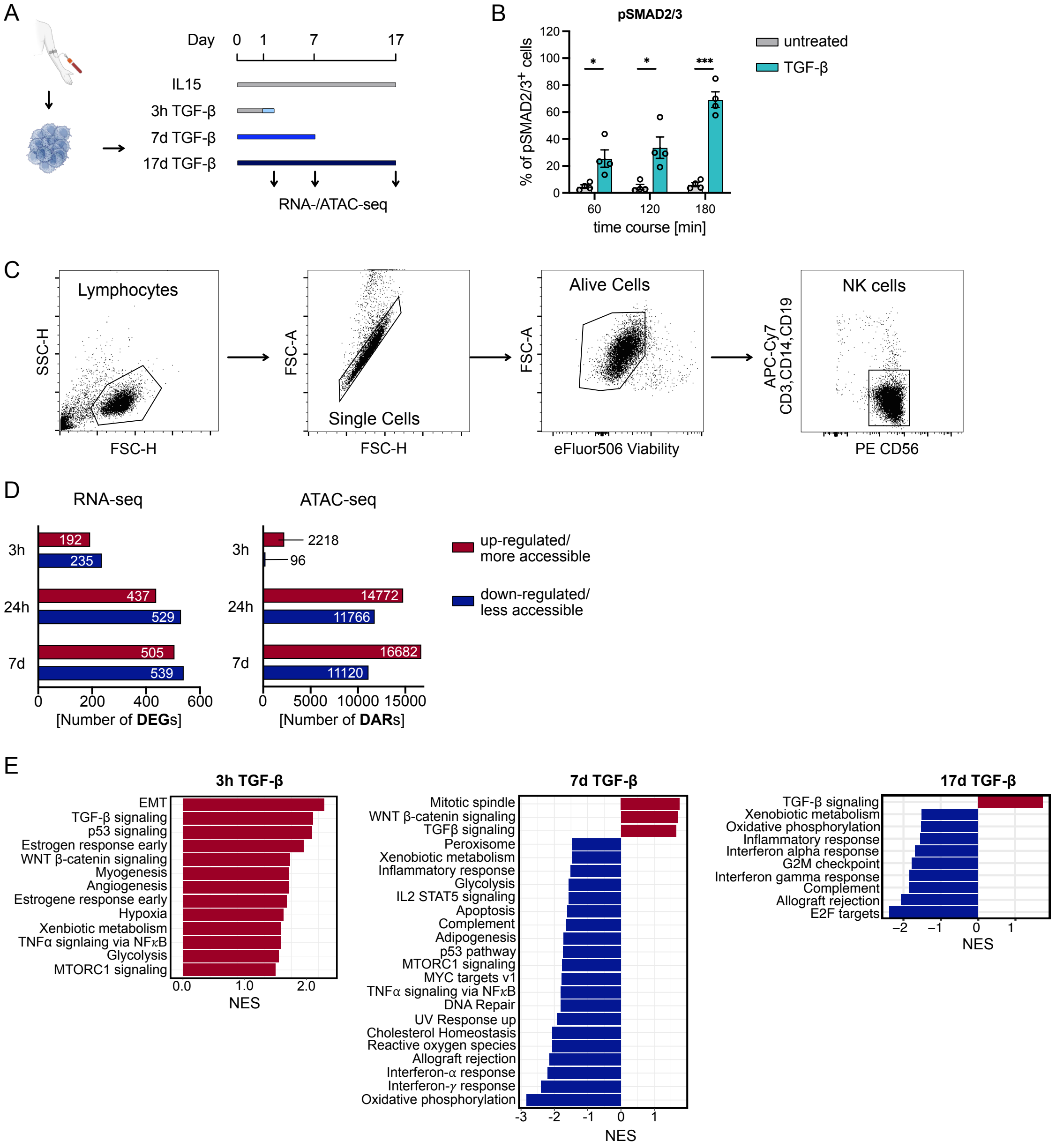


**Supplementary Figure 1: TGF-β-induced transcriptional and epigenetic changes. (A)** Experimental design. Enriched and purified peripheral NK cells from healthy donors were cultured with IL-15 (50 ng/mL) alone, IL-15 (50 ng/mL) plus TGF-β (10 ng/mL) for 7 d and 17 d, or IL-15 (50 ng/mL) for 1 d followed by 3 h TGF-β (10 ng/mL) stimulation on day 1. **(B)** Bar plot shows pSMAD2/3 levels of 7 d cultivated NK cells (50 ng/mL IL15) followed by a 3 h stimulation (10ng/mL TGF-β). Significance was calculated using a parametric paired student-t-test (*p < 0.05; **p < 0.01; ***p < 0.001; ****p < 0.0001). **(C)** Gating strategy for Lineage$^-$ CD56$^+$ NK cells**. (D)** Number of significant DARs and DEGs (padj < 0.05, |log2(FC)| > 0.58)) for 3 h, 7 d and 17 d (n = 5). Red indicates increased accessibility or expression; blue indicates decreased accessibility or expression. **(E)** GSEA (database: Hallmark) of DEGs (padj < 0.05), showing all significantly enriched pathways as normalised enrichments score (NES, padj(NES) < 0.05) for 3 h, 7 d and 17 d TGF-β treated NK cells.

Supplementary Figure 2

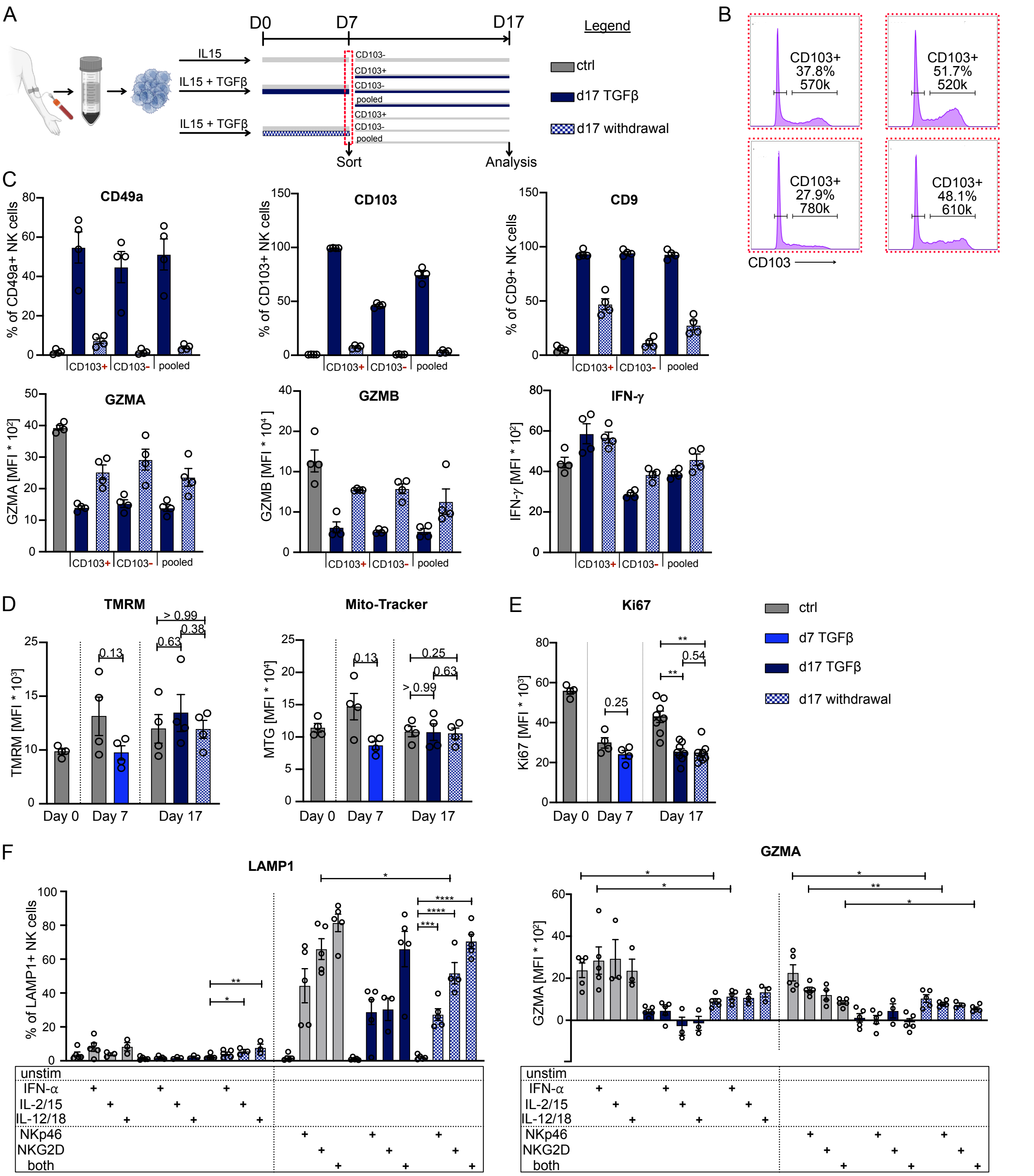


**Supplementary Figure 2: Analysis of long-term TGF-β-stimulated NK cells. (A)** Experimental setup. On day 7, cells were FACS-sorted based on CD103 expression. After sorting, CD103$^{high}$ and CD103$^{low}$ populations were normalized to the same cell number (based on the condition with the lowest yield) and cultured in fresh media with or without TGF-β for an additional 10 days. **(B)** Histogram shows CD103 sort gate. Cells were pre-gated on lymphocytes, single cells, live cells, lineage-negative, and CD56$^{+}$. Depicted are the frequency and total number of collected cells (n=4, one experiment). **(C)** Bar plots show frequency of CD49a$^+$, CD103$^+$, and CD9$^+$ cells and MFI for GZMA, GZMB, and IFN-γ on day 17 (n=4, one experiment). **(D)** Bar plots show expression levels of TMRM staining assessing mitochondrial membrane potential and MitoTracker Green (MTG) on day 17 (n=4, two independent experiments). **(E)** Bar plots show expression of Ki67 on day 1, day 7 (n=4), and day 17 (n=8) (three independent experiments). **(F)** Bar plots show frequency of LAMP1$^+$ NK cells after restimulation on day 17 (n=4, two independent experiments). Significance was calculated only between control and withdrawal conditions and within withdrawal. Only significant differences are shown. Across the entire figure, statistical significance was calculated using a Wilcoxon signed-ranked test for panels C, D, and E, and a paired Student-t-test for panel F (*p < 0.05; **p < 0.01; ***p < 0.001; ****p < 0.0001).

Supplementary Figure 3

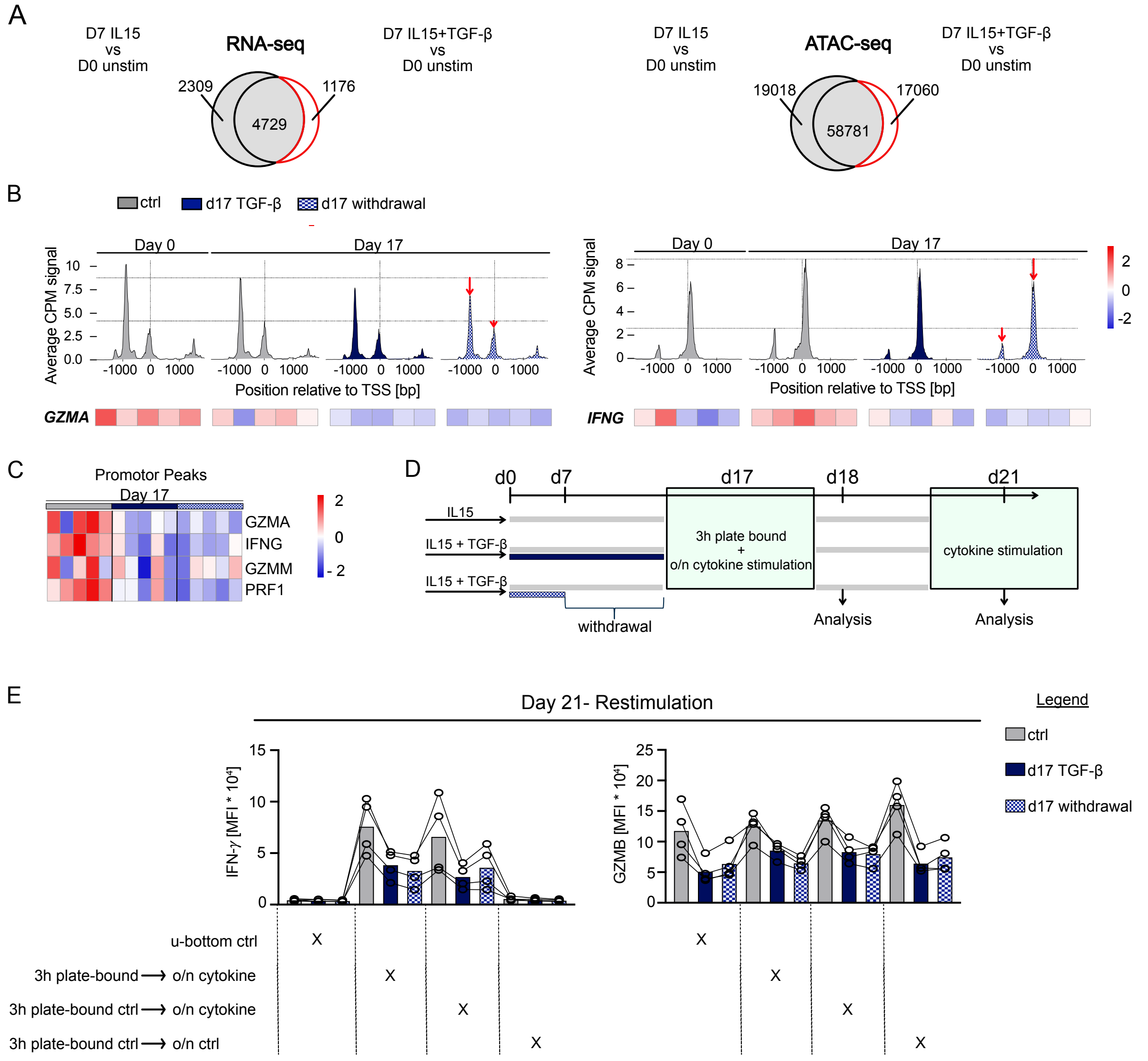


**Supplementary Figure 3: Transcriptional and epigenetic analysis of long-term TGF-β-stimulated NK cells. (A)** Venn diagram depicting the identification of TGF-β-regulated genes (|log2FC| > 0.58, padj < 0.05) by comparing DEGs in d 7 TGF-β vs unstimulated cells with d 7 IL-15 vs d 0 unstimulated cells. Genes uniquely regulated by TGF-β treatment (present in d 7 TGF-β vs d 0 unstimulated but absent in d 7 IL-15 vs d 0 unstimulated, marked by red line) were defined as TGF-β-regulated genes. **(B)** Line plots show average ATAC-seq signal (CPM) across the promoter region (± 2 kb from TSS) of the indicated genes, with signal extracted from BigWig files in 10 bp bins and averaged across biological replicates per condition. Vertical reference lines are aligned to the control condition. Arrows indicate the strength of reduced accessibility. Additionally, heatmap shows TSS peaks where each column represents one biological replicate. **(C)** Heatmap shows differentially accessible regions (DARs) annotated to transcription start sites (TSS) for effector gene loci, displayed as z-score of regularised counts. Each column depicts one biological replicate (n=5). **(D)** On day 17, NK cells were stimulated for 3 hours with anti-NKp46 and/or anti-NKG2D antibodies [10 ng/mL] or left unstimulated, followed by overnight culture in cytokine-supplemented medium (IL-12 [20 ng/mL], IL-18 [10 ng/mL], IL-2 [25 ng/mL], and IL-15 [50 ng/mL]). On day 18, cells were analysed by flow cytometry and transferred to fresh medium containing IL-15 [50 ng/mL] only for an additional 3 days. On day 21, cells were restimulated with IL-12 and IL-18 for 3 hours prior to final analysis. **(E)** Functional re-assessment of NK cells on day 21 following restimulation with IL-12 and IL-18 for 3 hours. IFN-γ and granzyme B (GZMB) expression are shown across four stimulation conditions applied on day 17. Conditions are indicated below each group. Colours represent day-17 treatment groups as in (E). Lines connect paired donors.

Supplementary Figure 4

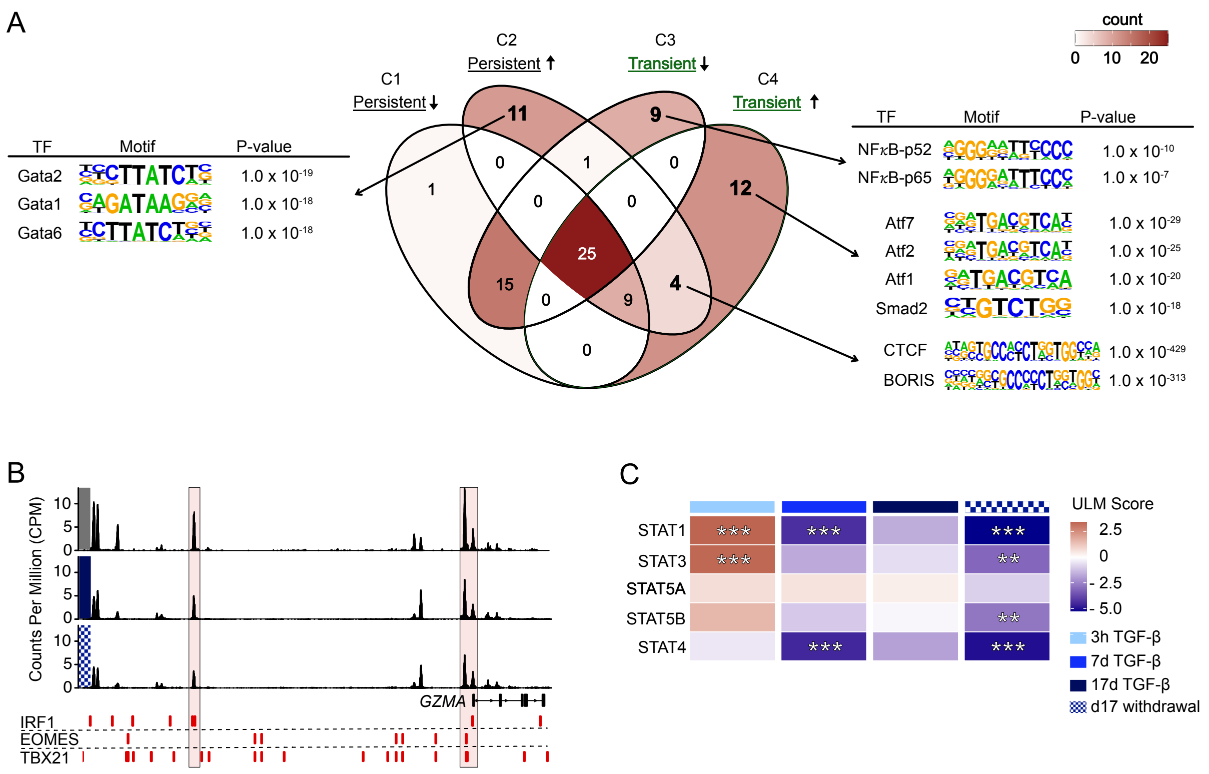


**Supplementary Figure 4: TF levels and activity after prolonged TGF-β treatment. (A)** TF motif enrichment analysis of ATAC-seq regions previous analysis (Fig.3D, C1, C2, C3, C4). The Venn diagram depicts the overlap of the top 50 enriched TF motifs (ranked by p value) across the indicated region categories; color intensity reflects the number of TFs assigned to each region. HOMER analysis was performed on pre-filtered regions (persistent down/up; transient down/up). For TFs present in more than one region, the most significant p value is shown. **(B)** Genome browser tracks of ATAC-seq signal at the GZMA locus from one representative donor, shown as normalized read coverage. Predicted TF binding sites within promoter and putative enhancer regions are indicated. Boxes highlight regions of interest. **(C)** Transcription factor (TF) activity was inferred using the Univariate Linear Model (ULM) method from the decoupleR framework, applied to differentially expressed genes (DEGs) from each pairwise condition comparison. ULM scores reflect the estimated activity of each TF based on the enrichment of its target genes within the DEG set. Only TFs with positive mode-of-regulation (MOR) interactions and more than 10 target genes in the TGFβ-regulated gene set were included. Significance of TF activity is indicated as * $p < 0.05$, ** $p < 0.01$, *** $p < 0.001$.

Supplementary Figure 5

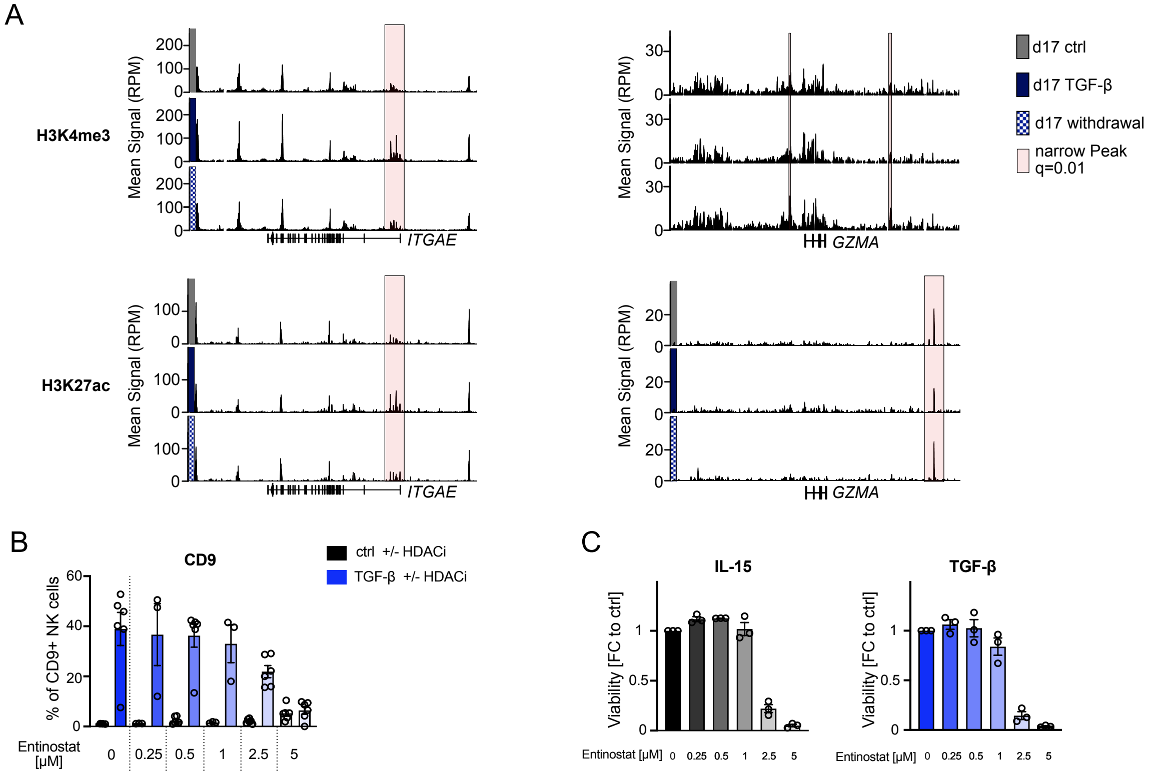


**Supplementary Figure 5: Analysis of permissive histone marks at TR and effector loci (A)** Mean CUT&RUN signal (RPM) for H3K4me3 (top) and H3K27ac (bottom) is shown at the ITGAE (left, endcoding CD103) and GZMA (right) loci for 17 d control (gray), 17 d TGF-β (dark blue), and 17 d withdrawal (blue hatched) conditions. Each track represents the mean signal across three biological replicates. Red shaded boxes indicate narrow peaks called at $q < 0.01$ (MACS3). Gene models are shown below the tracks. **(B)** Bar graphs show the frequency of CD9+ NK cells at d 7 (n=3-6; two independent experiments. Cells were treated with control (black) or TGF-β (blue) in the presence of increasing concentrations of Entinostat (0, 0.25, 0.5, 1, 2.5, 5 µM). Individual data points represent biological replicates; bars show mean ± SEM. **(C)** NK cell viability following treatment with increasing concentrations of the HDAC inhibitor Entinostat (0, 0.25, 0.5, 1, 2.5, and 5 µM) under IL-15 (left, grey) or TGF-β (right, blue) culture conditions. Viability is expressed as fold change (FC) relative to the untreated control (0 µM). Each dot represents an individual donor; bars indicate mean ± SEM.

Supplementary Figure 6

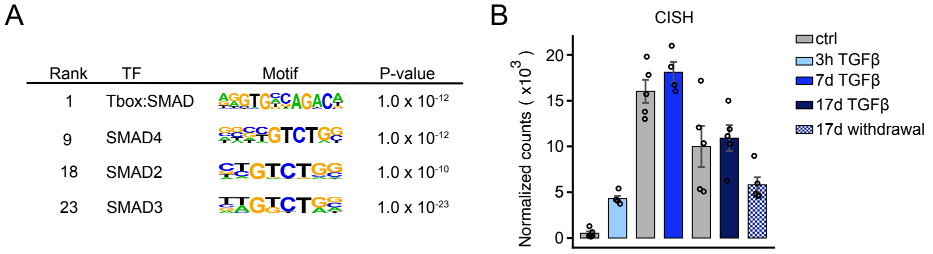


**Supplementary Figure 6: Analysis of SMAD-bound regions. (A)** HOMER analysis of SMAD-bound regions (n = 2774). Selected motifs are shown with their rank, associated TF, motif logo, and p-value. **(B)** Normalized RNA-seq counts for CISH across conditions: untreated control (ctrl, grey), 3h TGF-β (light blue), 7 d TGF-β (blue), 17 d TGF-β (dark blue), and 17 d TGF-β withdrawal (hatched). Individual data points represent biological replicates; bars show mean ± SEM.

Supplementary Figure 7

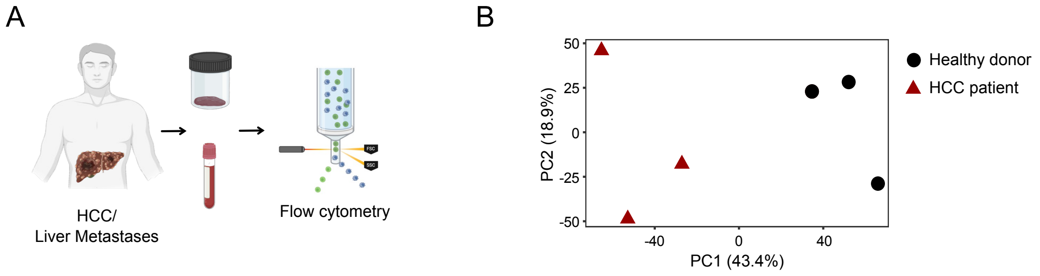


**Supplementary Figure 7: Analysis of patient NK cells (A)** Experimental setup. NK cells from human HCC tumors or liver metastases of other cancer entities were analyzed by flow cytometry (n = 10). In parallel, peripheral blood NK cells from HCC patients were enriched, cultured overnight with IL-15 (10 ng/mL) and IL-2 (25 U/mL), restimulated on day 1 with IL-12/IL-18 (50 ng/mL) for 4 h, and analyzed by flow cytometry. **(B)** ATAC-seq analysis of sorted peripheral NK cells from healthy donors and HCC patients with reduced NK-cell functionality (n = 3). Principal component analysis (PCA) shows variance along PC1 and PC2 based on the top 80,000 most variable features.
